# Dynamic tracking and identification of tissue-specific secretory proteins in the circulation of live mice

**DOI:** 10.1101/2020.09.21.299198

**Authors:** Kwang-eun Kim, Isaac Park, Jeesoo Kim, Myeong-Gyun Kang, Won Gun Choi, Hyemi Shin, Jong-Seo Kim, Hyun-Woo Rhee, Jae Myoung Suh

**Affiliations:** Graduate School of Medical Science and Engineering, KAIST, Daejeon, Republic of Korea; Department of Chemistry, Seoul National University, Seoul, Republic of Korea; Center for RNA Research, Institute for Basic Science, Seoul, Republic of Korea; School of Biological Sciences, Seoul National University, Seoul, Republic of Korea; Department of Chemistry, Ulsan National Institute of Science and Technology (UNIST), Ulsan, Republic of Korea

## Abstract

Here we describe *i*SLET (*<u>i</u>n situ* <u>S</u>ecretory protein <u>L</u>abeling via <u>E</u>R-anchored <u>T</u>urboID) which labels secretory pathway proteins as they transit through the ER-lumen to enable dynamic tracking of tissue-specific secreted proteomes *in vivo*. We expressed *i*SLET in the mouse liver and demonstrated efficient *in situ* labeling of the liver-specific secreted proteome which could be tracked and identified within circulating blood plasma. *i*SLET is a versatile and powerful tool for studying spatiotemporal dynamics of secretory proteins, a valuable class of biomarkers and therapeutic targets.

## Main text

Secretory proteins released into the blood circulation play essential roles in physiological systems and are core mediators of interorgan communication^1^. To investigate this critical class of proteins, previous studies analyze conditioned media from *in vitro* or *ex vivo* culture models to identify cell type-specific secretory proteins, but these models often fail to fully recapitulate the intricacies of multi-organ systems and thus do not sufficiently reflect *in vivo* realities^2^. In other approaches, bioinformatic tools such as QENIE (Quantitative Endocrine Network Interaction Estimation) have been developed^3^, however, *in silico* predictions of endocrine protein factors still require many additional layers of experimental validation. These limitations provide compelling motivation to develop *in vivo* techniques that can identify and resolve characteristics of tissue-specific secretory proteins along time and space dimensions.

To address this gap, we sought to utilize recently developed proximity-labeling enzymes such as engineered biotin ligase (BioID)^4^ or ascorbate peroxidase (APEX)^5^. When provided appropriate substrates, these enzymes generate reactive biotin species, leading to *in situ* biotinylation of proximal proteins on lysine or tyrosine residues, respectively. Thereafter, the biotinylated proteins are readily enriched through streptavidin affinity purification and can be identified through mass spectrometry. Recently, TurboID, a newly engineered biotin ligase, was developed to overcome the low biotinylation efficiency of BioID in subcellular compartments^6^. TurboID exhibits a 100-fold improvement in efficiency compared to that of BioID in the endoplasmic reticulum (ER) of cultured human cells^6^. Here, we introduce a new tool to profile tissue-specific secretory proteins by *in situ* proximity labeling of ER lumen proteins through the catalytic actions of an ER-anchored TurboID.

To engineer a TurboID based tool for labeling secretory proteins located in the ER lumen, we first tested the functionality of two ER lumen-targeted TurboIDs, an ER lumen-localized TurboID (TurboID-KDEL) and an ER membrane-anchored TurboID (Sec61b-TurboID), in cultured cells. We transfected either TurboID-KDEL or Sec61b-TurboID expression constructs, both of which also express a V5 epitope tag, to cultured mammalian cells and analyzed biotinylated proteins in cell lysates and culture supernatant (**Fig. 1a**). Immunofluorescence analysis of transfected cells with anti-V5 antibody and fluorescence-conjugated streptavidin confirmed expected patterns of ER localization for both TurboID-KDEL and Sec61b-TurboID along with their biotinylated targets **(Fig. 1b)**, consistent with results from previous ER localization studies for APEX2-KDEL and Sec61b-APEX2^7^. Analysis of biotinylated proteins in control cell lysates revealed the presence of several endogenous biotinylated carboxylases^8^ which were not detected in culture supernatant, indicating that these carboxylases are not secreted (**Fig. 1c**). In contrast to control cells, a broad array of biotinylated proteins was detected in both the cell lysate and culture supernatant of cells expressing TurboID-KDEL and Sec61b-TurboID in a biotin treatment-dependent manner (**Fig. 1c**).

**Fig. 1.**
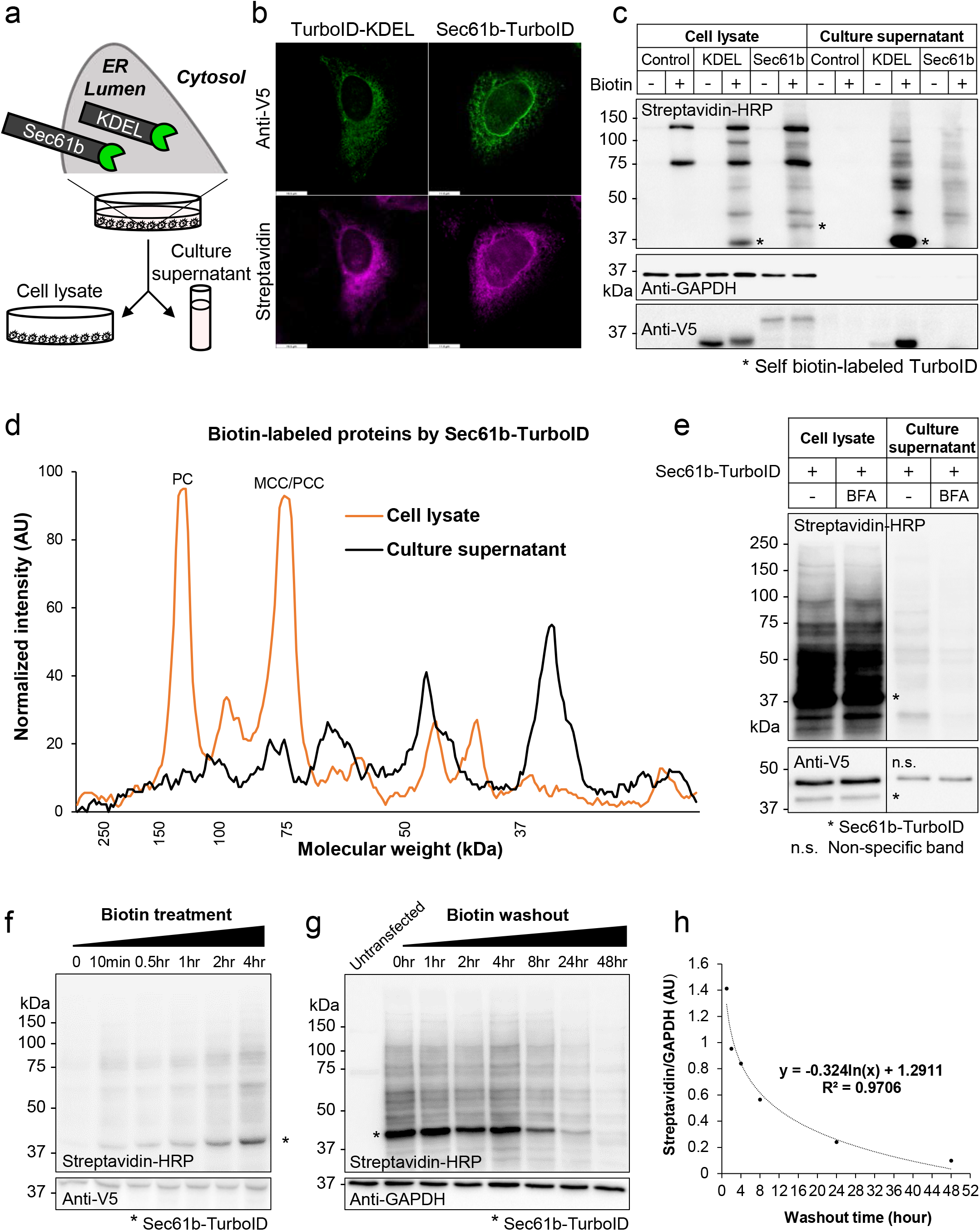
Proximity labeling of secretory pathway proteins using ER-anchored TurboID. **a**, Schematic illustration for secretory protein labeling by ER-localized TurboID (TurboID-KDEL) or ER-anchored TurboID (Sec61b-TurboID). **b**, Immunofluorescence localization of TurboID (Anti-V5) and biotinylated proteins (Streptavidin-Alexa) in HeLa cells transfected with TurboID-KDEL or Sec61b-TurboID expression plasmids. **c**, Western blots for biotinylated proteins (Streptavidin-HRP) and TurboID (Anti-V5) in cell lysates or culture supernatant of NIH-3T3 cells transfected with GFP (Control), TurboID-KDEL (KDEL), or Sec61b-TurboID (Sec61b) expression plasmids. Anti-GAPDH is a loading control. Asterisk indicates self-biotinylated TurboID-KDEL or Sec61b-TurboID. **d**, Line-scan analysis of biotinylated proteins in cell lysate (orange) or culture supernatant (black) from NIH-3T3 cells transfected with Sec61b-TurboID expression plasmids and treated with biotin. PC, Pyruvate carboxylase; MCC/PCC, Methylcrotonyl-CoA Carboxylase/ Propionyl-CoA carboxylase. **e**, Effect of Brefeldin A (BFA) on biotinylated protein secretion in HepG2 cells transfected with Sec61b-TurboID expression plasmids. **f**, Time course blot for biotinylated-labeled proteins (Streptavidin-HRP) in cell lysates of HepG2 cells transfected with Sec61b-TurboID expression plasmids. **g**, Time course blot for biotinylated protein (Streptavidin-HRP) turnover in cell lysates of HepG2 cells transfected with Sec61b-TurboID expression plasmids following biotin washout. **h**, Quantitation and plotting of the time course blot for biotinylated protein turnover shown in **g**. Asterisk indicates Sec61b-TurboID.

Somewhat unexpectedly, we found that TurboID-KDEL localization was not exclusive to the ER compartment and TurboID-KDEL itself was secreted and readily detectable in the culture supernatant of biotin treated cells (**Fig. 1c**). On the other hand, the ER-anchored Sec61b-TurboID was undetectable in the culture supernatant (**Fig. 1c** and **Supplementary Fig. 1**). These data indicate effective retention of Sec61b-TurboID, but not TurboID-KDEL, in the ER compartment through ER membrane-tethering action of the single transmembrane domain of Sec61b. We also confirmed that Sec61b-TurboID robustly labeled secretory proteins without self-secretion in a HepG2 human liver cell line, whereas TurboID-KDEL was again found to be secreted into the culture supernatant (**Supplementary Fig. 2**).

Notably, the pattern of biotinylated proteins generated by Sec61b-TurboID in the culture supernatant was clearly different from that of whole cell lysate, which is expected as ER-resident proteins and secretory proteins differ in composition **(Fig. 1d)**. To further confirm the secretory pathway origin of Sec61b-TurboID biotinylated proteins, we treated HepG2 cells expressing Sec61b-TurboID with Brefeldin A (BFA), an inhibitor of ER to Golgi protein transport, and observed a uniform reduction in the amount of biotinylated proteins detected in the culture supernatant **(Fig. 1e)**. Taken together, these data indicate that catalytically active Sec61b-TurboID is expressed and faithfully retained in the ER-lumen, a necessary property for *in vivo* applications that require efficient and accurate labeling of tissue-specific secretory proteins.

Labeling kinetics determined by biotin treatment time course studies indicate that Sec61b-TurboID efficiently labels secretory proteins in HepG2 cells by 10 min with increased labeling up to 4 hr (**Fig. 1f**). Conversely, biotin washout time course studies indicate that Sec61b-TurboID labeled secretory proteins are largely sustained for 8 hr (**Fig. 1g and 1h**). Therefore, Sec61b-TurboID can efficiently label classical secretory proteins in a biotin-dependent manner indicating compatibility with kinetic studies such as classical pulse-chase labeling analyses.

Next, we applied our method, named *i*SLET, *<u>i</u>n situ* <u>S</u>ecretory protein <u>L</u>abeling via <u>E</u>R-anchored <u>T</u>urboID, in live mice to demonstrate its *in vivo* functionality. Sec61b-TurboID adenovirus was delivered to mice via tail vein injection to establish a liver *i*SLET mouse model and biotin was administered to these mice to induce labeling of liver secretory proteins (**Fig. 2a**). Because TurboID has faster labeling kinetics than BioID, we administered 24 mg/kg biotin to mice for 3 days, as compared to 5 to 7 days of biotin administration used for previous BioID labeling studies in the postnatal brain^9,10^. Four days after Sec61b-TurboID adenovirus delivery, we observed that Sec61b-TurboID expression was restricted to the liver (**Supplementary Fig. 3**). Liver tissues examined by histological analysis did not reveal any obvious adverse effects due to adenoviral overexpression of TurboID and biotin administration (**Supplementary Fig. 3**).

**Fig. 2.**
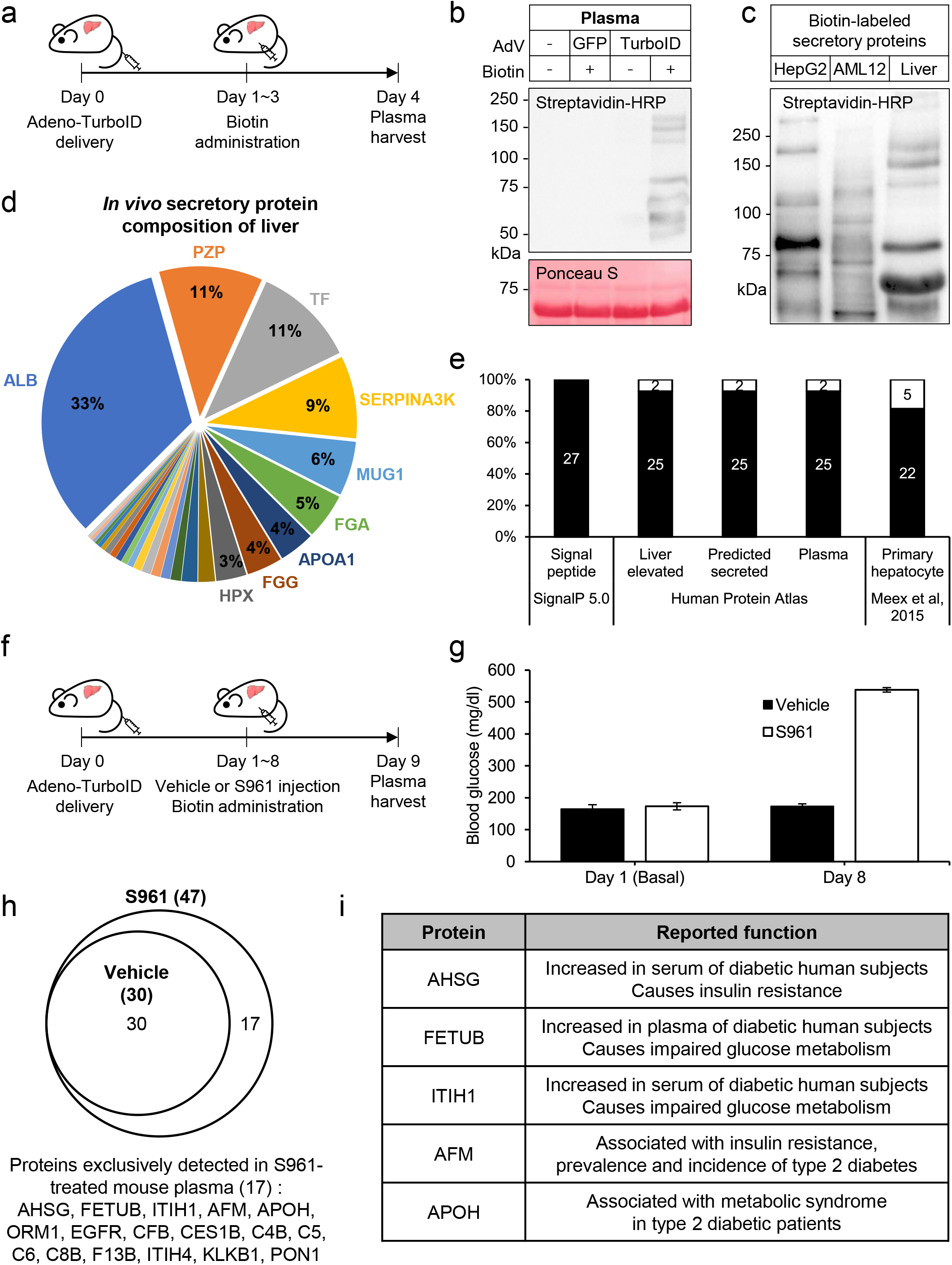
Identification of liver-specific secretory proteins in mouse plasma from liver *i* SLET mice. **a**, Experimental scheme for adenoviral expression of Sec61b-TurboID and biotin labeling in mouse liver tissue. **b**, Streptavidin-HRP detection of biotinylated proteins and Ponceau S detection of proteins from mouse plasma after adenoviral delivery of Sec61b-TurboID. **c**, Biotinylated secretory protein profiles generated by Sec61b-TurboID in supernatants of hepatocyte cell lines, HepG2 and AML12, and plasma of liver *i*SLET mice. **d**, Relative abundance of biotinylated secretory proteins detected in plasma of liver *i*SLET mice. ALB, Serum albumin; PZP, Pregnancy zone protein; TF, Serotransferrin; SERPINA3K, Serine protease inhibitor A3k; MUG1, Murinoglobulin-1; FGA, Fibrinogen alpha chain; APOA1, Apolipoprotein A-I; FGG, Fibrinogen gamma chain; HPX, Hemopexin. **e**, Specificity analysis for biotinylated proteins with SignalP 5.0, human protein atlas and literature. **f**, Experimental scheme for adenoviral expression of Sec61b-TurboID and biotin labeling in the S961-induced insulin resistance model. **g**, Blood glucose in Vehicle or S961 injected mouse. n=3 per group. **h**, Biotinylated secretory proteins detected in plasma of Vehicle or S961 injected liver *i*SLET mice. AHSG, Alpha-2-HS-glycoprotein; FETUB, Fetuin-B; ITIH1, Inter-alpha-trypsin inhibitor heavy chain H1; AFM, Afamin; APOH, Beta-2-glycoprotein 1; ORM1, Alpha-1-acid glycoprotein 1; EGFR, Receptor protein-tyrosine kinase; CFB, Complement factor B; CES1B, Carboxylic ester hydrolase; C4B, Complement C4-B; C5, Complement C5; C6, Complement component 6; C8B, Complement component C8 beta chain; F13B, Coagulation factor XIII B chain; ITIH4, Inter alpha-trypsin inhibitor, heavy chain 4; KLKB1, Plasma kallikrein; PON1, Serum paraoxonase/arylesterase 1. **i**. Representative candidates related with insulin resistance detected in this study.

As expected, and consistent with results obtained from the culture supernatant of Sec61b-TurboID-expressing cell lines, endogenous biotinylated proteins were not detected in plasma samples from liver *i*SLET mice (**Fig. 2b**). Thus, we could unambiguously detect TurboID-dependent biotinylated liver secretory proteins in the plasma without any background (**Fig. 2b**). Interestingly, the pattern of biotinylated proteins secreted from the liver *in vivo* was unique and clearly distinct from that of the secretory protein profile of hepatocyte cell lines, human HepG2 and mouse AML12 (**Fig. 2c**). These data confirm the *in vivo* functionality of Sec61b-TurboID in liver tissues as demonstrated by the detection of biotinylated secretory protein species in the plasma of liver *i*SLET mice.

We next performed proteomic analysis of biotinylated proteins enriched from liver *i*SLET mice plasma via liquid chromatography and tandem mass spectrometry (LC-MS/MS). Here, we followed a previously optimized mass spectrometric identification workflow^11,12^ which provides direct evidence for biotinylated peptides identified by the mass shift of the biotinylated lysine residue. From our LC-MS/MS data, 27 biotinylated proteins were identified in Sec61b-TurboID mouse plasma **(Fig. 2d and Supplementary Table 1)**. Representative MS/MS spectra of the biotinylated peptides from our optimized workflow show the accurate identification of biotinylated residues (**Supplementary Fig. 4 and Supplementary Table 2)**. Signal peptide analysis for the biotinylated proteins with SignalP 5.0^13^ revealed that all of the detected proteins contain signal peptides required for cotranslational transport to the ER-lumen **(Fig. 2e)**.

As expected, serum albumin (ALB) was the most abundant biotinylated protein detected from liver *i*SLET mice plasma samples **(Fig. 2d)**. Interestingly, the second most abundant protein was pregnancy zone protein (PZP, Q61838) (**Fig. 2d)**, which is also annotated under the alias alpha-2-macroglobulin (A2M, Q6GQT1) in the UniProt database. However, *Pzp* and *A2m* are independent genes in the mouse genome^14^, and the identified peptides in our analysis were a precise match to the sequence of PZP but not A2M **(Supplementary Fig. 4)**.

We found that 93% of the proteins identified in this study are annotated as liver-enriched and predicted as secreted plasma proteins in the Human Protein Atlas database **(Fig. 2e)**. We next compared the secretory protein profiles from liver *i*SLET mice plasma with *ex vivo* secretome studies using primary hepatocytes^15^. While a considerable fraction (81%) of proteins were common in both (**Fig. 2e**), fibrinogen gamma chain (FGA), complement component C8 alpha chain (C8A), histidine-rich glycoprotein (HRG), inter alpha-trypsin inhibitor, heavy chain 4 (ITIH4) and serine protease inhibitor A3M (SERPINA3M) were only detected in mouse plasma of liver *i*SLET mice (**Supplementary Table. 1)**. Taken together, our results indicate that the liver-specific secretory protein profiles obtained from liver *i*SLET mice are conserved in human and more accurately reflect *in vivo* physiology compared to conventional *ex vivo* secretome analyses.

We next applied *i*SLET to characterize secreted proteomes associated with *in vivo* pathophysiology in which endocrine signals play an important role such as insulin resistance. S961 is an insulin receptor antagonist that induces systemic insulin resistance^16^. We administered S961 and biotin for 8 consecutive days to liver *i*SLET mice generated by adenoviral delivery of the Sec61-TurboID transgene (**Fig. 2f**). S691 administration to mice dramatically increased blood glucose confirming the insulin resistance state (**Fig. 2g**). Proteomic analysis of biotinylated proteins from vehicle (PBS) or S961 group plasma identified 30 and 47 protein species, respectively (**Fig. 2h**). Notably, 17 of the identified proteins were exclusively found in the S961 administered insulin resistant group. Among these proteins, many have been reported to play a role in the development of insulin resistance (**Fig. 2i**).

Alpha-2-HS-glycoprotein (AHSG), also known as Fetuin-A, is elevated in serum of obese diabetic human subjects and induces insulin resistance^17^. Fetuin-B (FETUB) is increased in type 2 diabetes patients and causes glucose intolerance^15^. Inter-alpha-trypsin inhibitor heavy chain H1 (ITIH1) is increased in human subjects with impaired glucose tolerance or diabetes and antibody neutralization of ITIH1 ameliorates systemic insulin resistance in mice^18^. Afamin (AFM) is strongly associated with insulin resistance, prevalence and incidence of type 2 diabetes in pooled analysis in >20,000 individuals^19^. Beta-2-glycoprotein 1 (APOH), also known as apolipoprotein H is increased in the plasma of type 2 diabetic patients^20^. These results demonstrate that *i*SLET technology can be successfully applied to animal disease models for the discovery of tissue-specific secreted proteins with potential value as therapeutic targets or biomarkers.

*i*SLET is the first application of proximity labeling to dynamically track tissue-specific secretory proteins in the circulation of live mice. Liver *i*SLET mice may be utilized to deepen our understanding of liver endocrine signaling by investigating secretory protein profiles under various physiological or disease conditions. Another valuable feature of *i*SLET technology is that it can be applied to longitudinal secretome profiling studies by drawing blood samples, which contain labeled secretome, at multiple time points from the same individual. Pre-immunodepletion of abundant plasma proteins such as ALB and PZP can further enhance coverage of secretory protein profiles identified from *i*SLET studies.

Furthermore, *i*SLET is a versatile and adaptable *in vivo* approach to profile tissue-specific secretory proteins as *i*SLET expression in a tissue-of-interest can be achieved using a variety of existing conditional gene expression strategies^21^. *i*SLET will be a valuable experimental tool for the identification of tissue-specific endocrine proteins and the deconvolution of complex interorgan communication networks.

## Methods

### Animals

All animal experiments were approved by the KAIST institutional animal care and use committee. 10 week old C57BL/6J (JAX, 000664) male mice were used for all animal experiments.

Mice were maintained under a 12 h light-dark cycle in a climate-controlled specific pathogen-free facility within the KAIST Laboratory Animal Resource Center. Standard chow diet (Envigo, 2018S) and water were provided *ad libitum*. Tissues were dissected and fixed for histological analysis or snap-frozen in liquid nitrogen until further analysis.

### Cell culture and transfection

All cell lines were purchased from the American Type Culture Collection (ATCC; www.atcc.org) and cultured according to standard mammalian tissue culture protocols at 37℃, 5% CO_2_ in a humidified incubator. NIH-3T3 cells were cultured in DMEM (Hyclone, SH30243.01) supplemented with 10% bovine serum (Invitrogen, 16170-078) and antibiotics (100 units/mL penicillin, 100 μg/mL streptomycin). HepG2 cells were cultured in DMEM (Hyclone, SH30243.01) supplemented with 10% fetal bovine serum (Gibco, 16000-044), 1% GlutaMax (Gibco, 35050061) and antibiotics (100 units/mL penicillin, 100 μg/mL streptomycin). AML12 cells were cultured in DMEM/F12 (Gibco, 11320-033) supplemented with 10% FBS, 1% Insulin-Transferrin-Selenium (Gibco, 41400-045) and antibiotics. 293AD cells and HeLa cells were cultured in DMEM supplemented with 10% FBS and antibiotics. For transient plasmid transfection, cells were plated at 2.5×10^5^ cells/well in a 6-well culture plate. 24 h after plating, cells were transfected using 6 μL jetPEI (Polyplus) and 2.5 μg GFP, TurboID-KDEL, or Sec61b-TurboID plasmids according to manufacturer protocols.

### *In vitro* biotin labeling and cell lysate preparation

5 mM Biotin (Sigma, B4639) stock was prepared in DPBS with NaOH titration. 24 h after plasmid transfection or adenoviral transduction, cells were washed with PBS and further maintained for 16 hr in culture medium supplemented with 50 μM biotin. For the biotin washout experiment, following biotin labeling, cells were washed with PBS and further maintained in fresh culture medium. Cells were lysed by RIPA (Pierce, 89901) with Xpert Protease Inhibitor Cocktail (GenDEPOT, P3100-010) and incubated 30 min at 4℃. Lysates were cleared by centrifugation at 16,000 g for 20 min at 4℃. The clear supernatant was used for western blots. Protein concentrations were determined by BCA assay (Pierce, 23225).

### Culture supernatant protein preparation

Cells were washed with PBS twice and the culture medium was changed to phenol red free DMEM (Hyclone, SH30284.01) supplemented 1mM pyruvate (Sigma, S8636) with or without 50 μM biotin. For secretory pathway inhibition, 1X GolgiPlug™ (BD, 555029), which contains Brefeldin A, was treated with biotin. 16 h after biotin incubation, culture supernatant was centrifuged at 400 g for 5 min and the supernatant was filtered by 0.22 μm PES syringe filter (Millipore, SLGP033RB). The filtered supernatant was concentrated by Amicon Ultra 2 mL 10K (Millipore, UFC201024) with buffer exchange to 50 mM Tris-HCl pH 6.8. Concentrated supernatant was used for western blots. Protein concentrations were determined by BCA assay.

### Western blots

Denatured proteins were separated on 12% SDS-PAGE gels. Separated proteins were transferred to PVDF membrane (Immobilon-P, IPVH00010). Membranes were stained with Ponceau S for 15 min, washed with TBS-T (25 mM Tris, 150 mM NaCl, 0.1% Tween 20, pH 7.5) twice for 5 min, and photographed. Membranes were blocked in 3% BSA in TBS-T for 1 h, washed with TBS-T five times for 5 min each and incubated with primary antibodies, Anti-V5 (Invitrogen, R960-25, 1:10000), Anti-GAPDH (CST, 14C10, 1:5000), in 3% BSA in TBS-T for 16 h at 4℃. Then, membranes were washed five times with TBS-T for 5 min each and incubated with secondary anti-mouse antibodies (Vector, PI-2000, 1:10000) or anti-rabbit antibodies (Vector, PI-1000, 1:10000) for 1 h at room temperature. For detecting biotinylated proteins, blocked membranes were incubated with streptavidin-HRP (Thermo, 21126, 1:15000) in 3% BSA in TBS-T for 1 h at room temperature. Membranes were washed five times in TBS-T before detection with chemiluminescent HRP substrate (Immobilon, P90720) and imaged on a ChemiDoc™ XRS+ system (Bio-Rad, 1708265).

### Immunofluorescence staining

HeLa cells were plated on round coverslips (thickness no. 1, 18 mm radius) and transfected with plasmids. Cells were treated with 50 μM biotin for 30 min. Cells were fixed with 4% paraformaldehyde and permeabilized with ice-cold methanol for 5 min at −20°C. Next, cells were washed with DPBS and blocked for 1 h with 2% dialyzed BSA in DPBS at room temperature. Cells were incubated 1 h at room temperature with the primary antibody, Anti-V5 (Invitrogen, R960-25, 1:5000), in blocking solution. After washing four times with TBS-T each 5 min, cells were simultaneously incubated with secondary Alexa Fluor 488 goat anti-mouse immunoglobulin G (IgG) (Invitrogen, A-11001, 1:1000) and Streptavidin-Alexa Fluor 647 IgG (Invitrogen, S11226, 1:1000) for 30 min at room temperature. Cells were then washed four times with TBS-T each 5 min. Immunofluorescence images were obtained and analyzed using a Confocal Laser Scanning Microscope (Leica, SP8X) with White Light Laser (WLL): 470 – 670 nm (1 nm tunable laser) and HyD detector.

### Adenovirus production and infection

Recombinant adenoviruses were generated as previously described^22^. Briefly, Sec61b-TurboID was cloned to the pAdTrack-CMV shuttle vector by KpnI and NotI digestion. The cloned shuttle vector was linearized with PmeI and transformed to BJ5183-AD-1 cells. The recombinant adenoviral plasmid was linearized with PacI and transfected to 293AD cells. Stepwise amplification of adenovirus was performed, and adenovirus was concentrated by ViraBind™ adenovirus purification kit (Cell Biolabs, VPK-100). Adenovirus titer was measured by counting GFP-positive cells 24 h after infection with serial dilution. For adenoviral infection, cells were plated at 2.5 × 10^5^ cells/well in a 6-well culture plate. 24 h after plating, cells were infected with 1.25 × 10^6^ adenoviral GFP or Sec61b-TurboID particles.

### *In vivo* biotin labeling and protein sample preparation

Approximately 10^8^ adenoviral GFP or Sec61b-TurboID particles were injected to mice via the tail vein. 24 mg/ml biotin stock was prepared in DMSO. Vehicle (10% DMSO in PBS) or Biotin solution (2.4 mg/mL) was filtered through a 0.22 μm PES syringe filter and injected 10 μL/g (24 mg/kg) by daily intraperitoneal injection for 3 consecutive days. For the acute insulin resistance model, S961 (100 nmol/kg, Novo Nordisk) was delivered by daily intraperitoneal injection for 8 consecutive days, 2 hours prior to daily biotin injection. Biotin was not administered on the last day to minimize residual biotin in blood. Blood samples were obtained by cardiac puncture and plasma was separated in BD Microtainer® blood collection tubes (BD, 365985). Tissues were lysed and homogenized in RIPA buffer with Xpert Protease Inhibitor Cocktail (GenDEPOT, P3100-010) by FastPrep-24™ bead homogenizer (MP Biomedicals). Lysates were clarified by three rounds of centrifugation at 16,000 g for 20 min at 4℃ and supernatant collection. The clear supernatant was used for western blots. Protein concentrations were determined by BCA assay.

### Peptide sample preparation and enrichment of biotinylated peptides

Plasma samples were first subjected to buffer exchange with PBS to completely remove residual free biotin via 10k MWCO filtration for three times. The biotin depleted plasma samples were transferred and denatured with 500 μL of 8 M urea in 50 mM ammonium bicarbonate for 1 h at 37°C, and followed by reduction of disulfide bonds with 10 mM dithiothreitol for 1 h at 37°C. The reduced thiol groups in the protein samples were subsequently alkylated with 40 mM iodoacetamide for 1 h at 37°C in the dark. The resulting alkylated samples were diluted eight times using 50 mM ABC and subjected to trypsinization at 2% (w/w) trypsin concentration under 1 mM CaCl_2_ concentration for overnight in Thermomixer (37°C and 500 rpm). Samples were centrifuged at 10,000 g for 3 min to remove insoluble material. Then, 150 μL of streptavidin beads (Pierce, 88816) per replicate was washed with 2 M urea in TBS four times and combined with the individual digested sample. The combined samples were rotated for 1 h at room temperature. The flow-through fraction was kept, and the beads were washed twice with 2 M urea in 50 mM ABC and finally with pure water in new tubes. The bound biotinylated peptides were eluted with 400 μL of 80% acetonitrile containing 0.2 % TFA and 0.1 % formic acid after mixing and heating the bead slurry at 60°C. Each eluate was collected into a new tube. The elution process was repeated four more times. Combined elution fractions were dried using Vacufuge® (Eppendorf) and reconstituted with 10 μL of 25 mM ABC for further analysis by LC-MS/MS.

### LC-MS/MS analysis of enriched biotinylated peptides

The enriched samples were analyzed with an Orbitrap Fusion Lumos mass spectrometer (Thermo Scientific) coupled with a NanoAcquity UPLC system (Waters, Milford) in sensitive acquisition settings. Precursor ions were acquired at a range of m/z 400–1600 with 120 K resolving power and the isolation of precursor for MS/MS analysis was performed with a 1.4 Th. Higher-energy collisional dissociation (HCD) with 30% collision energy was used for sequencing with a target value of 1e5 ions determined by automatic gain control. Resolving power for acquired MS2 spectra was set to 30k at m/z 200 with 150 ms maximum injection time. The peptide samples were loaded onto the trap column (3 cm × 150 µm i.d) via the back-flushing technique and separated with a 100 cm long analytical capillary column (75 µm i.d.) packed in-house with 3 µm Jupiter C18 particles (Phenomenex, Torrance). The long analytical column was placed in a dedicated 95 cm long column heater (Analytical Sales and Services) regulated to a temperature of 45°C. NanoAcquity UPLC system was operated at a flow rate of 300 nL/min over 2 h with a linear gradient ranging from 95% solvent A (H2O with 0.1% formic acid) to 40% of solvent B (acetonitrile with 0.1% formic acid).

### LC-MS/MS data processing and the identification of biotinylated peptides

All MS/MS datasets were first subject to peak picking and mass recalibration processed with RawConverter^23^ (http://fields.scripps.edu/rawconv) and MZRefinery^24^ (https://omics.pnl.gov/software/mzrefinery) software, respectively, and then were searched by MS-GF+^25^ algorithm (v.9979) at 10 ppm precursor ion mass tolerance against the UniProt reference proteome database (55,152 entries, Mouse). The following search parameters were applied: semi-tryptic digestion, fixed carbamidomethylation on cysteine, dynamic oxidation of methionine, and dynamic biotinylation of a lysine residue (delta monoisotopic mass: +226.07759 Da). The False discovery rate (FDR) was set at < 0.5% for non-redundantly labeled peptide level and the resulting protein FDR was near or less than 1%. MS/MS spectrum annotation for biotinylated peptides was carried out using LcMsSpectator software (https://omics.pnl.gov/software/lcmsspectator).

### Histological analysis

Mouse livers were fixed in 10 % neutral buffered formalin (Sigma, HT501128) for 24 hr and embedded in paraffin by an automated tissue processor (Leica, TP1020). 4 μm-thick tissue sections were obtained, deparaffinized, rehydrated, and stained with hematoxylin and eosin.

## Acknowledgments

We thank Dr. K.-J. Oh for assistance in adenovirus generation, Dr. K. Kim for assistance in mass spectrometry analysis and H.-S. Jung, J.E. Kim, and H. Jung for technical support. We thank all lab members for helpful discussions and technical assistance. This work was supported by the National Research Foundation (NRF) of Korea (NRF-2018R1A2A3075389, NRF-2016M3A9B6902871) and Global Research Laboratory (GRL) Program through the NRF funded by the Ministry of Science and ICT (No. NRF-2017K1A1A2013124). J.-S.K. thanks to the support from Institute for Basic Science from the Ministry of Science and ICT of Korea (IBS-R008-D1).

## Author contributions

K.K., I.P., J.-S.K., H.-W.R. and J.M.S. designed research, K.K., I.P., J.K., M.K., W.G.C., and H.S. conducted research. K.K., I.P., J.-S.K., H.-W.R. and J.M.S. wrote the manuscript.

