## Supplementary file for "Dynamic tracking and identification of tissue-specific secretory proteins in the circulation of live mice"

### Kim et al., Supplementary Figure 1

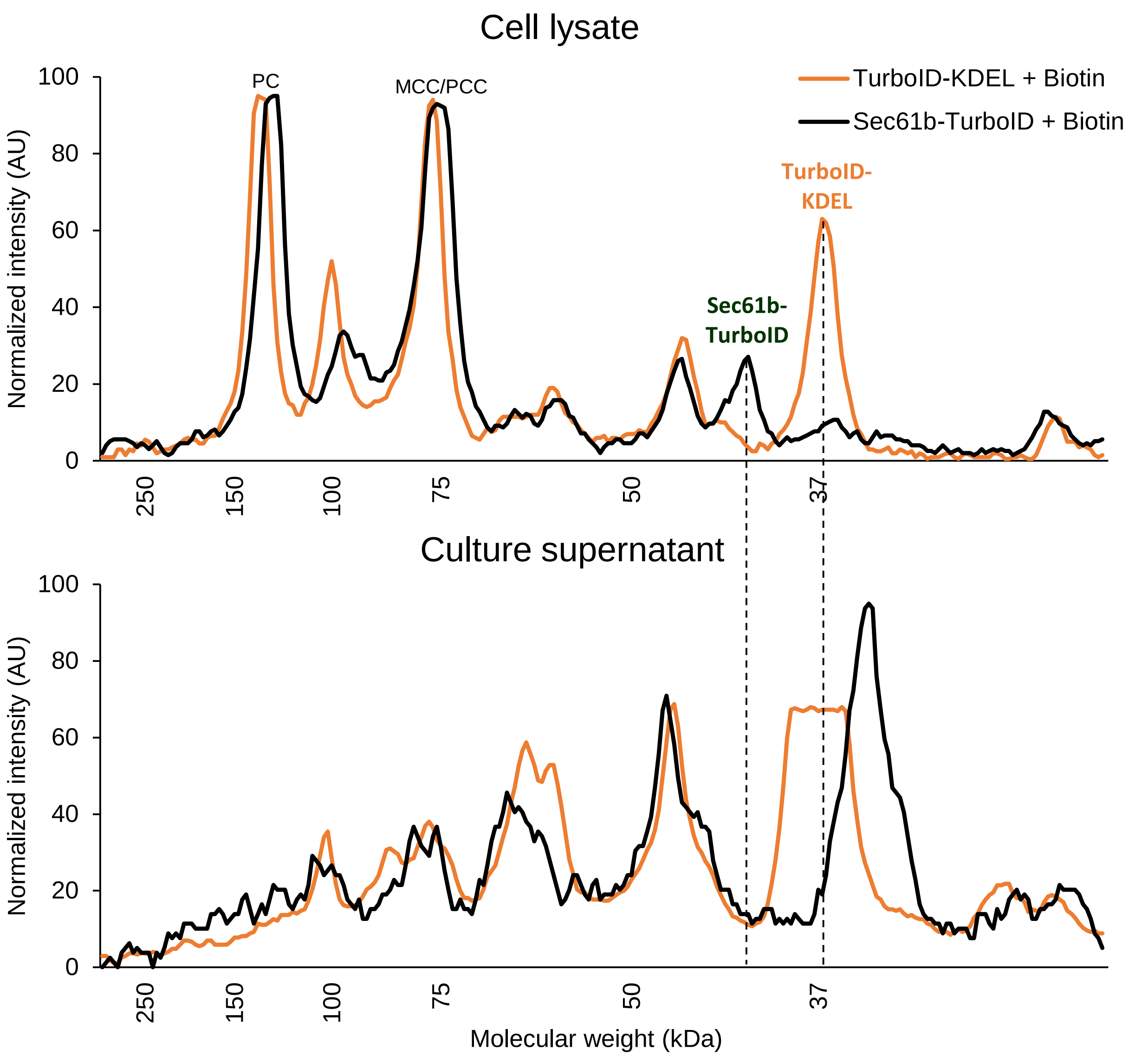


**Supplementary Fig. 1 Biotin-labeled protein profiles of cell lysates and culture supernatants from TurboID-KDEL or Sec61b-TurboID expressing cells.** Line-scan analysis of biotin-labeled proteins detected by HRP-streptavidin in cell lysates and culture supernatants of NIH-3T3 cells transfected with TurboID-KDEL (orange) or Sec61b-TurboID (black) expression plasmids and treated with biotin. PC, Pyruvate carboxylase; MCC/PCC, Methylcrotonyl-CoA carboxylase/Propionyl-CoA carboxylase.

### Kim et al., Supplementary Figure 2

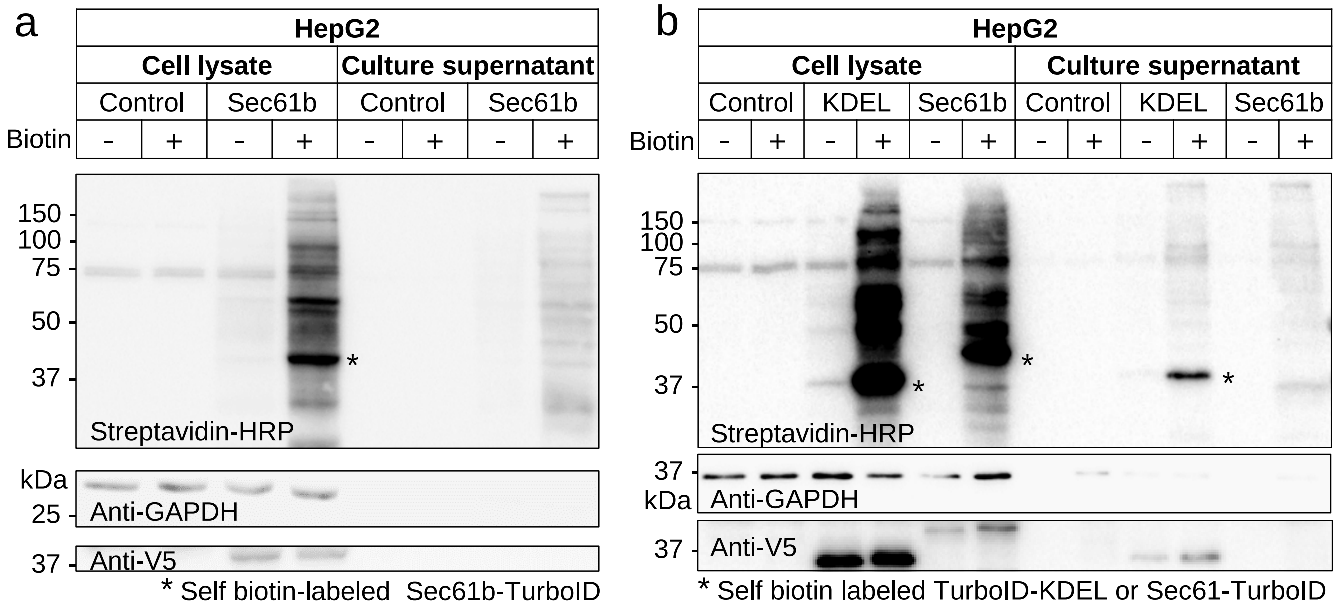


**Supplementary Fig. 2 Self-secretion of TurboID-KDEL in HepG2 cells. a**, Western blots for biotin-labeled proteins (Streptavidin-HRP) and Sec61b-TurboID (Anti-V5) in cell lysates or culture supernatants of HepG2 cells transfected with GFP (Control) or Sec61b-TurboID (Sec61b) expression plasmids. **b**, Western blots for biotin-labeled proteins (Streptavidin-HRP) and TurboID (Anti-V5) in cell lysates or culture supernatants of HepG2 cells transfected with GFP (Control), TurboID-KDEL (KDEL) or Sec61b-TurboID (Sec61b) expression plasmids. Anti-GAPDH is a loading control. Asterisk indicates self biotin-labeled TurboID-KDEL or Sec61-TurboID.

#### Kim et al., Supplementary Figure 3

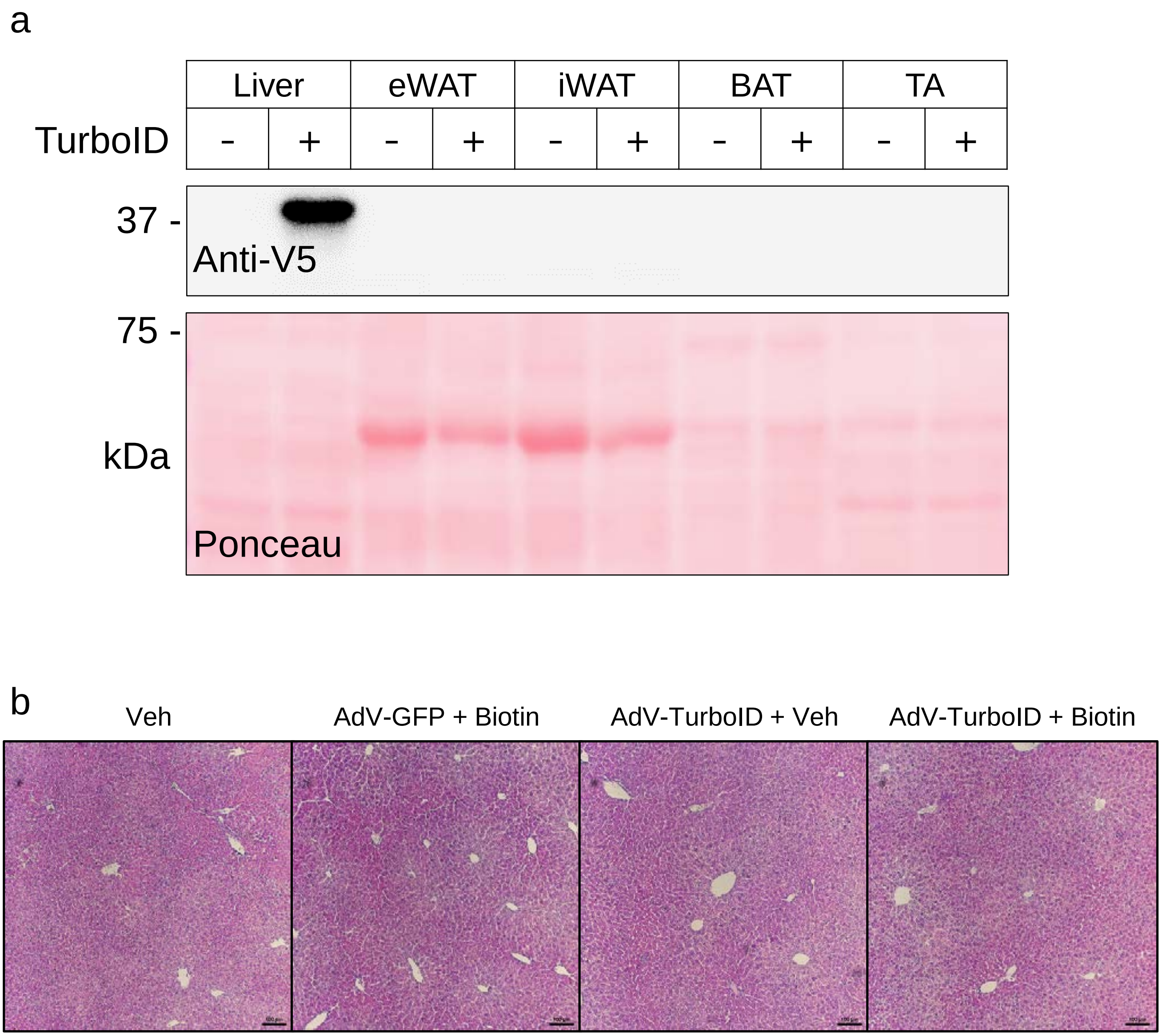


**Supplementary Fig. 3 Validation of liver-specific Sec61-TurboID expression and liver histology. a**, Western blot for Sec61b-TurboID (Ant-V5) in liver and other tissues. eWAT: epididymal White Adipose Tissue, iWAT: inguinal White Adipose Tissue, BAT: Brown Adipose Tissue, TA: Tibialis Anterior muscle. **b**, Hematoxylin and Eosin staining of liver tissues from mice with indicated treatment. Vehicle only (Veh), GFP expressing adenovirus (AdV-GFP), Sec61b-TurboID expressing adenovirus (AdV-TurboID).

### Kim et al., Supplementary Figure 4

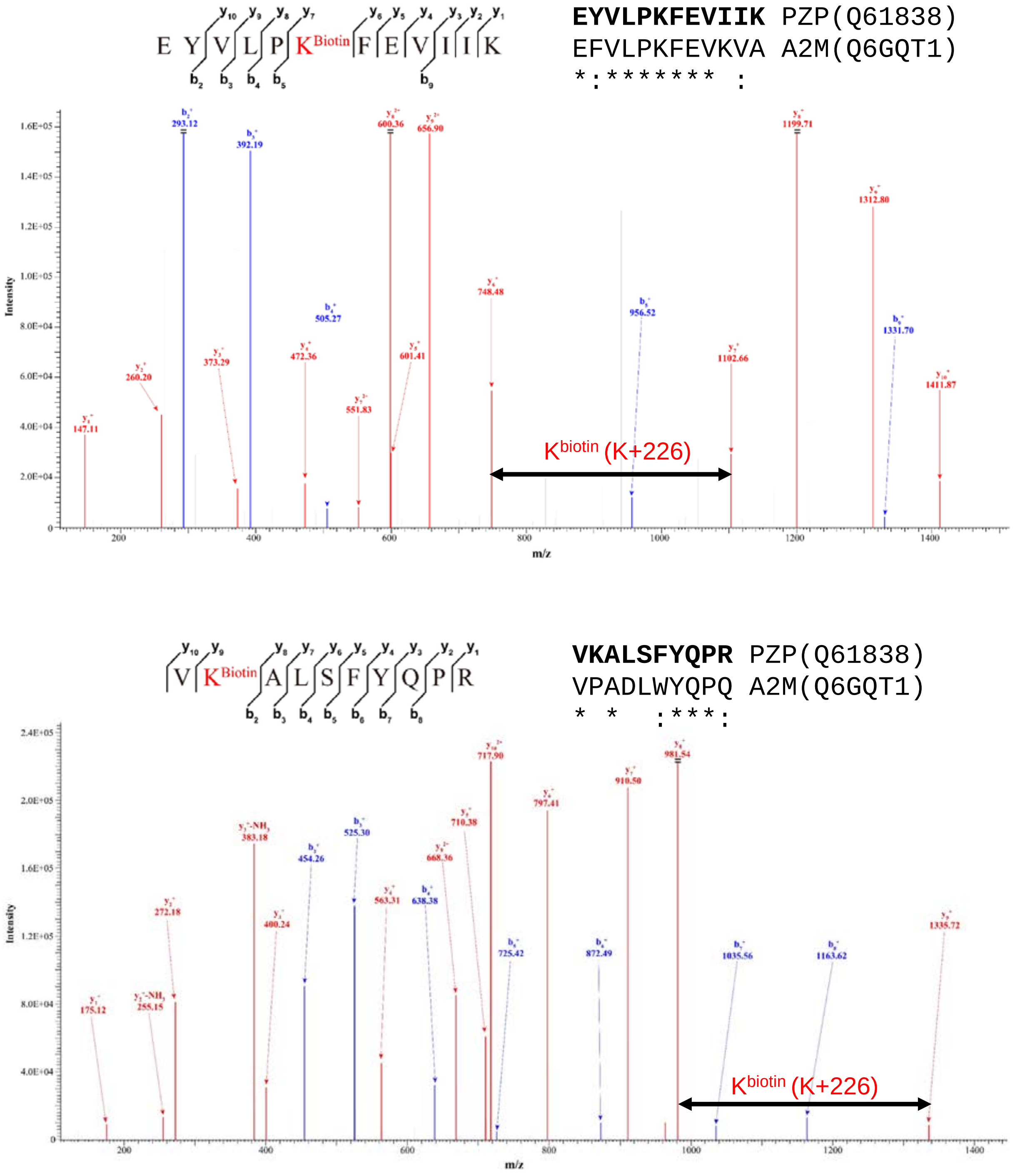


**Supplementary Fig. 4 Representative tandem mass spectrum of biotinylated PZP peptides.** Mass of the biotinylated lysine residue is 354 Da (K+226 Da). Arrow represents the mass shift of the biotinylated lysine residue from the biotinylated PZP peptide (Q61838).

**Supplementary Table 1. Identified liver secretory proteins in mouse plasma with definitive *in situ* biotinylation evidence on lysine residues and comparison with other studies**

| | | Protein species retrieved by UniProt Retrieve/ID mapping | Adeno-Sec61b-TurboID+Biotin (Replicate 1) | | Adeno-Sec61b-TurboID+Biotin (Replicate 2) | | SignalP 5.0 (Armenteros et al, 2019) | Human Protein Atlas (www.proteinatlas.org) | | | Ex vivo secretome of primary hepatocyte (Meex et al, 2015) |
|---|---|---|---|---|---|---|---|---|---|---|---|
| UniProt accession | Gene name | Protein name (Both detected, non-redundant peptide >1) | Non-redundant peptide | PSM | Non-redundant peptide | PSM | Signal Peptide (Sec/SPI) Likelihood | Elevated expression in Liver? | Predicted secreted protein? | Plasma protein? | Peptide count |
| P07724 | Alb | Albumin | 43 | 134 | 45 | 161 | 0.993 | Yes | Yes | Yes | 145 |
| Q61838 | Pzp | Pregnancy zone protein | 20 | 43 | 21 | 56 | 0.997 | Yes | Yes | Yes | 6 |
| Q921I1 | Tf | Serotransferrin | 21 | 42 | 28 | 56 | 0.998 | Yes | Yes | Yes | 75 |
| A0A0R4J0I1 | Serpina3k | Serine protease inhibitor A3K | 16 | 46 | 13 | 33 | 0.981 | Yes (SERPINA3) | Yes (SERPINA3) | Yes (SERPINA3) | 21 |
| P28665 | Mug1 | Murinoglobulin-1 | 13 | 19 | 24 | 33 | 0.949 | - | - | - | 6 |
| E9PV24 | Fga | Fibrinogen alpha chain | 12 | 18 | 16 | 26 | 0.995 | Yes | Yes | Yes | Not detected |
| Q00623 | Apoa1 | Apolipoprotein A-I | 7 | 13 | 10 | 21 | 0.997 | Yes | Yes | Yes | 9 |
| Q8VCM7 | Fgg | Fibrinogen gamma chain | 9 | 15 | 11 | 19 | 0.999 | Yes | Yes | Yes | 7 |
| Q91X72 | Hpx | Hemopexin | 7 | 15 | 7 | 14 | 0.988 | Yes | Yes | Yes | 17 |
| P22599 | Serpina1b | Alpha-1-antitrypsin 1-2 | 3 | 3 | 9 | 13 | 0.988 | Yes (SERPINA1) | Yes (SERPINA1) | Yes (SERPINA1) | 23 |
| P01027 | C3 | Complement C3 | 5 | 6 | 6 | 9 | 0.988 | Yes | Yes | Yes | 36 |
| Q61147 | Cp | Ceruloplasmin | 3 | 4 | 5 | 6 | 0.994 | Yes | Yes | Yes | 29 |
| O08677 | Kng1 | Kininogen-1 | 3 | 4 | 3 | 5 | 0.999 | Yes | Yes | Yes | 17 |
| Q61646 | Hp | Haptoglobin | 2 | 2 | 3 | 7 | 0.997 | Yes | Yes | Yes | 22 |
| Q8K0E8 | Fgb | Fibrinogen beta chain | 4 | 4 | 4 | 5 | 0.999 | Yes | Yes | Yes | 12 |
| P21614 | Gc | Vitamin D-binding protein | 2 | 3 | 3 | 5 | 0.996 | Yes | Yes | Yes | 40 |
| P19221 | F2 | Prothrombin | 2 | 2 | 3 | 4 | 0.900 | Yes | Yes | Yes | 11 |
| P20918 | Plg | Plasminogen | 2 | 2 | 4 | 4 | 0.993 | Yes | Yes | Yes | 1 |
| P23953 | Ces1c | Carboxylesterase 1C | 2 | 3 | 2 | 3 | 0.979 | - | - | - | 10 |
| P32261 | Serpinc1 | Antithrombin-III | 2 | 3 | 2 | 3 | 0.988 | Yes | Yes | Yes | 6 |
| Q8K182 | C8a | Complement component C8 alpha chain | 2 | 2 | 2 | 4 | 0.997 | Yes | Yes | Yes | Not detected |
| P06909 | Cfh | Complement factor H | 1 | 1 | 3 | 3 | 0.999 | Yes | Yes | Yes | 1 |
| Q9ESB3 | Hrg | Histidine-rich glycoprotein | 1 | 2 | 2 | 2 | 0.998 | Yes | Yes | Yes | Not detected |
| A6X935 | Itih4 | Inter alpha-trypsin inhibitor, heavy chain 4 | 1 | 1 | 2 | 2 | 0.960 | Yes | Yes | Yes | Not detected |
| Q03734 | Serpina3m | Serine protease inhibitor A3M | 2 | 2 | 1 | 1 | 0.967 | Yes (SERPINA3) | Yes (SERPINA3) | Yes (SERPINA3) | Not detected |
| Q61247 | Serpinf2 | Alpha-2-antiplasmin | 1 | 1 | 2 | 2 | 0.979 | Yes | Yes | Yes | 5 |
| Q61703 | Itih2 | Inter-alpha-trypsin inhibitor heavy chain H2 | 1 | 1 | 2 | 2 | 0.998 | Yes | Yes | Yes | 16 |

**Supplementary Table 2. Identified peptide-to-spectrum (PSM) match list of biotinylated peptides in mouse plasma.**

**Biotinylated peptides are highlighted**

| #SpecFile | ScanNum | FragMethod | Precursor | IsotopeError | PrecursorError(ppm) | Charge | Pre | Peptide | Post | Protein | DeNovoScore | MSGFScore | SpecEValue | EValue |
|---|---|---|---|---|---|---|---|---|---|---|---|---|---|---|
| Issac_4_1_NAQ444_J100_190802_FIXED_rere.mzML | 28338 | HCD | 821.079 | 0 | 3.64 | 3 | K | QSLVQATEENLNK+226.078AVNYASR | I | sp\|A6X935\|ITIH4_MOUSE | 146 | 129 | 8.89E-19 | 2.24E-11 |
| Issac_4_1_NAQ444_J100_190802_FIXED_rere.mzML | 6020 | HCD | 539.594 | 0 | -0.57 | 3 | K | EK+226.078VTSSGTSTTHR | S | sp\|E9PV24\|FIBA_MOUSE | 123 | 123 | 2.75E-17 | 6.64E-10 |
| Issac_4_1_NAQ444_J100_190802_FIXED_rere.mzML | 8794 | HCD | 687.825 | 0 | 0.27 | 2 | R | NNK+226.078DSNSLTR | N | sp\|E9PV24\|FIBA_MOUSE | 80 | 79 | 3.09E-12 | 7.28E-05 |
| Issac_4_1_NAQ444_J100_190802_FIXED_rere.mzML | 9524 | HCD | 575.783 | 0 | -1.48 | 2 | R | SCSK+226.078TITK | T | sp\|E9PV24\|FIBA_MOUSE | 64 | 61 | 5.49E-11 | 1.27E-03 |
| Issac_4_1_NAQ444_J100_190802_FIXED_rere.mzML | 17545 | HCD | 599.319 | 0 | 1.12 | 2 | K | ELLPTK+226.078DR | Q | sp\|E9PV24\|FIBA_MOUSE | 87 | 82 | 1.16E-10 | 2.67E-03 |
| Issac_4_1_NAQ444_J100_190802_FIXED_rere.mzML | 19146 | HCD | 665.311 | 0 | 3.58 | 3 | R | M(ox)K+226.078GLIDEANQDFTNR | I | sp\|E9PV24\|FIBA_MOUSE | 116 | 72 | 4.59E-12 | 1.12E-04 |
| Issac_4_1_NAQ444_J100_190802_FIXED_rere.mzML | 19212 | HCD | 997.461 | 0 | 2.20 | 2 | R | M(ox)K+226.078GLIDEANQDFTNR | I | sp\|E9PV24\|FIBA_MOUSE | 144 | 116 | 1.61E-15 | 3.94E-08 |
| Issac_4_1_NAQ444_J100_190802_FIXED_rere.mzML | 19825 | HCD | 784.050 | 1 | 0.34 | 3 | R | M(ox)K+226.078GLIDEANQDFTNRINK | L | sp\|E9PV24\|FIBA_MOUSE | 125 | 97 | 3.51E-15 | 8.74E-08 |
| Issac_4_1_NAQ444_J100_190802_FIXED_rere.mzML | 19856 | HCD | 783.717 | 0 | 1.17 | 3 | R | M(ox)K+226.078GLIDEANQDFTNRINK | L | sp\|E9PV24\|FIBA_MOUSE | 116 | 83 | 2.67E-14 | 6.64E-07 |
| Issac_4_1_NAQ444_J100_190802_FIXED_rere.mzML | 21189 | HCD | 870.112 | 0 | 1.54 | 3 | R | EINLQDYEGHQK+226.078QLQQVIAK | E | sp\|E9PV24\|FIBA_MOUSE | 224 | 204 | 1.21E-24 | 3.06E-17 |
| Issac_4_1_NAQ444_J100_190802_FIXED_rere.mzML | 25787 | HCD | 702.730 | 0 | 0.61 | 3 | K | QLQQVIAK+226.078ELLPTKDR | Q | sp\|E9PV24\|FIBA_MOUSE | 201 | 201 | 1.00E-19 | 2.47E-12 |
| Issac_4_1_NAQ444_J100_190802_FIXED_rere.mzML | 26073 | HCD | 702.729 | 0 | -0.17 | 3 | K | QLQQVIAK+226.078ELLPTKDR | Q | sp\|E9PV24\|FIBA_MOUSE | 115 | 94 | 1.79E-13 | 4.41E-06 |
| Issac_4_1_NAQ444_J100_190802_FIXED_rere.mzML | 26290 | HCD | 747.383 | 0 | 0.73 | 2 | K | LK+226.078NSLFDFQR | N | sp\|E9PV24\|FIBA_MOUSE | 123 | 120 | 1.45E-13 | 3.42E-06 |
| Issac_4_1_NAQ444_J100_190802_FIXED_rere.mzML | 27772 | HCD | 612.354 | 0 | 0.80 | 3 | K | QLQQVIAK+226.078ELLPTK | D | sp\|E9PV24\|FIBA_MOUSE | 130 | 128 | 1.17E-14 | 2.84E-07 |
| Issac_4_1_NAQ444_J100_190802_FIXED_rere.mzML | 27781 | HCD | 918.027 | 0 | 0.80 | 2 | K | QLQQVIAK+226.078ELLPTK | D | sp\|E9PV24\|FIBA_MOUSE | 176 | 168 | 9.23E-17 | 2.25E-09 |
| Issac_4_1_NAQ444_J100_190802_FIXED_rere.mzML | 30470 | HCD | 970.813 | 1 | 0.02 | 3 | K | M(ox)SPVPDLVPGSFK+226.078SQLQEAPPEWK | A | sp\|E9PV24\|FIBA_MOUSE | 219 | 147 | 6.89E-16 | 1.77E-08 |
| Issac_4_1_NAQ444_J100_190802_FIXED_rere.mzML | 30506 | HCD | 970.479 | 0 | 0.06 | 3 | K | M(ox)SPVPDLVPGSFK+226.078SQLQEAPPEWK | A | sp\|E9PV24\|FIBA_MOUSE | 196 | 136 | 1.53E-17 | 3.93E-10 |
| Issac_4_1_NAQ444_J100_190802_FIXED_rere.mzML | 31834 | HCD | 810.426 | 0 | 0.60 | 3 | R | QYLPALK+226.078M(ox)SPVPDLVPGSFK | S | sp\|E9PV24\|FIBA_MOUSE | 177 | 168 | 1.20E-23 | 3.02E-16 |
| Issac_4_1_NAQ444_J100_190802_FIXED_rere.mzML | 32160 | HCD | 810.426 | 0 | 1.21 | 3 | R | QYLPALK+226.078M(ox)SPVPDLVPGSFK | S | sp\|E9PV24\|FIBA_MOUSE | 92 | 27 | 1.46E-11 | 3.68E-04 |
| Issac_4_1_NAQ444_J100_190802_FIXED_rere.mzML | 20072 | HCD | 632.378 | 0 | -0.29 | 2 | K | IAPSK+226.078APILK | A | sp\|O08677\|KNG1_MOUSE | 104 | 75 | 2.62E-11 | 6.18E-04 |
| Issac_4_1_NAQ444_J100_190802_FIXED_rere.mzML | 31823 | HCD | 874.117 | 0 | 0.00 | 3 | R | VIEGTKTDGSPTFYSFK+226.078YLIK | E | sp\|O08677\|KNG1_MOUSE | 140 | 101 | 3.40E-14 | 8.60E-07 |
| Issac_4_1_NAQ444_J100_190802_FIXED_rere.mzML | 31866 | HCD | 874.452 | 1 | 0.51 | 3 | R | VIEGTKTDGSPTFYSFK+226.078YLIK | E | sp\|O08677\|KNG1_MOUSE | 108 | 70 | 1.65E-12 | 4.18E-05 |
| Issac_4_1_NAQ444_J100_190802_FIXED_rere.mzML | 36375 | HCD | 727.058 | 0 | 1.09 | 3 | K | ATSQVVAGTK+226.078YVIEFIAR | E | sp\|O08677\|KNG1_MOUSE | 108 | 57 | 4.88E-11 | 1.22E-03 |
| Issac_4_1_NAQ444_J100_190802_FIXED_rere.mzML | 32295 | HCD | 973.532 | 0 | 0.31 | 2 | K | FGPK+226.078VLNSINVAVFSK | K | sp\|O89020\|AFAM_MOUSE | 166 | 157 | 8.03E-18 | 1.98E-10 |
| Issac_4_1_NAQ444_J100_190802_FIXED_rere.mzML | 21262 | HCD | 708.381 | 0 | 0.52 | 3 | R | QVLTSEK+226.078TVLTGASGHLR | S | sp\|P01027\|CO3_MOUSE | 208 | 208 | 2.45E-22 | 6.11E-15 |
| Issac_4_1_NAQ444_J100_190802_FIXED_rere.mzML | 21432 | HCD | 708.715 | 1 | -0.40 | 3 | R | QVLTSEK+226.078TVLTGASGHLR | S | sp\|P01027\|CO3_MOUSE | 121 | 101 | 2.33E-17 | 5.81E-10 |
| Issac_4_1_NAQ444_J100_190802_FIXED_rere.mzML | 21458 | HCD | 871.929 | 0 | -3.36 | 2 | K | DSCIGTLVVK+226.078GDPR | D | sp\|P01027\|CO3_MOUSE | 102 | 91 | 1.20E-14 | 2.92E-07 |
| Issac_4_1_NAQ444_J100_190802_FIXED_rere.mzML | 27666 | HCD | 789.720 | 0 | -0.31 | 3 | R | AFYEHAPK+226.078QIFSAEFEVK | E | sp\|P01027\|CO3_MOUSE | 204 | 108 | 9.25E-13 | 2.31E-05 |
| Issac_4_1_NAQ444_J100_190802_FIXED_rere.mzML | 28102 | HCD | 958.793 | 0 | 3.18 | 3 | K | NYAGVFM(ox)DAGLAFK+226.078TSQGLQTEQR | A | sp\|P01027\|CO3_MOUSE | 103 | 45 | 1.77E-12 | 4.56E-05 |
| Issac_4_1_NAQ444_J100_190802_FIXED_rere.mzML | 28786 | HCD | 638.374 | 1 | -2.55 | 3 | R | VGLVAVDK+226.078GVFVLNKK | N | sp\|P01027\|CO3_MOUSE | 134 | 107 | 6.93E-15 | 1.71E-07 |
| Issac_4_1_NAQ444_J100_190802_FIXED_rere.mzML | 41416 | HCD | 686.702 | 0 | -0.62 | 3 | R | GVFDLGDAVSK+226.078ILQIEK | E | sp\|P01029\|CO4B_MOUSE | 94 | 58 | 3.69E-11 | 9.14E-04 |
| Issac_4_1_NAQ444_J100_190802_FIXED_rere.mzML | 34655 | HCD | 734.725 | 0 | 3.90 | 3 | R | CLTNLIEK+226.078VASYGVRPR | Y | sp\|P04186\|CFAB_MOUSE | 179 | 122 | 1.17E-13 | 2.89E-06 |
| Issac_4_1_NAQ444_J100_190802_FIXED_rere.mzML | 21911 | HCD | 855.912 | 0 | 1.14 | 2 | R | EK+226.078TSNFNADFALK | F | sp\|P06683\|CO9_MOUSE | 134 | 111 | 2.35E-14 | 5.68E-07 |
| Issac_4_1_NAQ444_J100_190802_FIXED_rere.mzML | 18271 | HCD | 613.619 | 0 | 1.99 | 3 | R | ITCYM(ox)GK+226.078WSTPPR | C | sp\|P06909\|CFAH_MOUSE | 80 | 42 | 1.30E-10 | 3.15E-03 |
| Issac_4_1_NAQ444_J100_190802_FIXED_rere.mzML | 887 | HCD | 629.257 | 0 | -1.84 | 2 | R | M(ox)K+226.078CSSM(ox)QK | F | sp\|P07724\|ALBU_MOUSE | 71 | 71 | 3.92E-12 | 9.04E-05 |
| Issac_4_1_NAQ444_J100_190802_FIXED_rere.mzML | 1017 | HCD | 629.258 | 0 | -0.48 | 2 | R | M(ox)K+226.078CSSM(ox)QK | F | sp\|P07724\|ALBU_MOUSE | 80 | 80 | 3.92E-12 | 9.04E-05 |
| Issac_4_1_NAQ444_J100_190802_FIXED_rere.mzML | 1159 | HCD | 629.258 | 0 | -0.10 | 2 | R | M(ox)K+226.078CSSM(ox)QK | F | sp\|P07724\|ALBU_MOUSE | 84 | 84 | 3.92E-12 | 9.04E-05 |
| Issac_4_1_NAQ444_J100_190802_FIXED_rere.mzML | 1317 | HCD | 629.259 | 0 | 0.48 | 2 | R | M(ox)K+226.078CSSM(ox)QK | F | sp\|P07724\|ALBU_MOUSE | 80 | 79 | 3.93E-12 | 9.07E-05 |
| Issac_4_1_NAQ444_J100_190802_FIXED_rere.mzML | 2975 | HCD | 629.259 | 0 | 0.19 | 2 | R | M(ox)K+226.078CSSM(ox)QK | F | sp\|P07724\|ALBU_MOUSE | 84 | 84 | 3.92E-12 | 9.04E-05 |
| Issac_4_1_NAQ444_J100_190802_FIXED_rere.mzML | 3235 | HCD | 629.259 | 0 | 0.39 | 2 | R | M(ox)K+226.078CSSM(ox)QK | F | sp\|P07724\|ALBU_MOUSE | 82 | 81 | 3.94E-12 | 9.10E-05 |
| Issac_4_1_NAQ444_J100_190802_FIXED_rere.mzML | 3498 | HCD | 629.258 | 0 | 0.00 | 2 | R | M(ox)K+226.078CSSM(ox)QK | F | sp\|P07724\|ALBU_MOUSE | 86 | 71 | 9.69E-12 | 2.23E-04 |
| Issac_4_1_NAQ444_J100_190802_FIXED_rere.mzML | 3763 | HCD | 629.259 | 0 | 0.19 | 2 | R | M(ox)K+226.078CSSM(ox)QK | F | sp\|P07724\|ALBU_MOUSE | 85 | 85 | 3.92E-12 | 9.04E-05 |
| Issac_4_1_NAQ444_J100_190802_FIXED_rere.mzML | 4036 | HCD | 629.259 | 0 | 0.19 | 2 | R | M(ox)K+226.078CSSM(ox)QK | F | sp\|P07724\|ALBU_MOUSE | 69 | 60 | 6.50E-11 | 1.50E-03 |
| Issac_4_1_NAQ444_J100_190802_FIXED_rere.mzML | 4286 | HCD | 629.258 | 0 | -0.29 | 2 | R | M(ox)K+226.078CSSM(ox)QK | F | sp\|P07724\|ALBU_MOUSE | 89 | 89 | 3.92E-12 | 9.04E-05 |
| Issac_4_1_NAQ444_J100_190802_FIXED_rere.mzML | 4348 | HCD | 621.261 | 0 | 0.39 | 2 | R | M(ox)K+226.078CSSMQK | F | sp\|P07724\|ALBU_MOUSE | 62 | 62 | 7.84E-12 | 1.81E-04 |
| Issac_4_1_NAQ444_J100_190802_FIXED_rere.mzML | 4539 | HCD | 629.258 | 0 | -0.10 | 2 | R | M(ox)K+226.078CSSM(ox)QK | F | sp\|P07724\|ALBU_MOUSE | 78 | 78 | 3.92E-12 | 9.04E-05 |
| Issac_4_1_NAQ444_J100_190802_FIXED_rere.mzML | 5316 | HCD | 621.261 | 0 | -0.59 | 2 | R | MK+226.078CSSM(ox)QK | F | sp\|P07724\|ALBU_MOUSE | 84 | 84 | 1.48E-12 | 3.42E-05 |
| Issac_4_1_NAQ444_J100_190802_FIXED_rere.mzML | 5338 | HCD | 629.259 | 0 | 0.39 | 2 | R | M(ox)K+226.078CSSM(ox)QK | F | sp\|P07724\|ALBU_MOUSE | 87 | 74 | 8.75E-12 | 2.02E-04 |
| Issac_4_1_NAQ444_J100_190802_FIXED_rere.mzML | 6275 | HCD | 468.567 | 0 | 0.26 | 3 | R | EAHK+226.078SEIAHR | Y | sp\|P07724\|ALBU_MOUSE | 120 | 116 | 1.75E-13 | 4.13E-06 |
| Issac_4_1_NAQ444_J100_190802_FIXED_rere.mzML | 6334 | HCD | 621.261 | 0 | 0.20 | 2 | R | M(ox)K+226.078CSSMQK | F | sp\|P07724\|ALBU_MOUSE | 75 | 71 | 7.97E-12 | 1.84E-04 |
| Issac_4_1_NAQ444_J100_190802_FIXED_rere.mzML | 11849 | HCD | 736.311 | 0 | 1.41 | 2 | K | CSSM(ox)QK+226.078FGER | A | sp\|P07724\|ALBU_MOUSE | 119 | 119 | 1.06E-14 | 2.51E-07 |
| Issac_4_1_NAQ444_J100_190802_FIXED_rere.mzML | 12123 | HCD | 736.310 | 0 | 0.33 | 2 | K | CSSM(ox)QK+226.078FGER | A | sp\|P07724\|ALBU_MOUSE | 66 | 53 | 7.76E-11 | 1.83E-03 |
| Issac_4_1_NAQ444_J100_190802_FIXED_rere.mzML | 12239 | HCD | 895.406 | 0 | 1.30 | 2 | R | VGTK+226.078CCTLPEDQR | L | sp\|P07724\|ALBU_MOUSE | 104 | 92 | 3.79E-14 | 9.17E-07 |
| Issac_4_1_NAQ444_J100_190802_FIXED_rere.mzML | 14233 | HCD | 758.023 | 0 | 0.56 | 3 | K | TPVSEHVTK+226.078CCSGSLVER | R | sp\|P07724\|ALBU_MOUSE | 165 | 165 | 1.59E-23 | 3.96E-16 |
| Issac_4_1_NAQ444_J100_190802_FIXED_rere.mzML | 15023 | HCD | 577.299 | 0 | 1.27 | 3 | K | LQTCCDKPLLK+226.078K | A | sp\|P07724\|ALBU_MOUSE | 147 | 143 | 2.63E-16 | 6.32E-09 |
| Issac_4_1_NAQ444_J100_190802_FIXED_rere.mzML | 15965 | HCD | 577.298 | 0 | 0.11 | 3 | K | LQTCCDK+226.078PLLKK | A | sp\|P07724\|ALBU_MOUSE | 139 | 139 | 2.49E-16 | 5.97E-09 |
| Issac_4_1_NAQ444_J100_190802_FIXED_rere.mzML | 16027 | HCD | 865.444 | 0 | 0.92 | 2 | K | LQTCCDK+226.078PLLKK | A | sp\|P07724\|ALBU_MOUSE | 136 | 118 | 3.32E-15 | 7.96E-08 |
| Issac_4_1_NAQ444_J100_190802_FIXED_rere.mzML | 16224 | HCD | 577.634 | 1 | 1.73 | 3 | K | LQTCCDK+226.078PLLKK | A | sp\|P07724\|ALBU_MOUSE | 102 | 80 | 4.17E-13 | 1.00E-05 |
| Issac_4_1_NAQ444_J100_190802_FIXED_rere.mzML | 16249 | HCD | 577.298 | 0 | -0.32 | 3 | K | LQTCCDK+226.078PLLKK | A | sp\|P07724\|ALBU_MOUSE | 82 | 68 | 3.59E-12 | 8.61E-05 |
| Issac_4_1_NAQ444_J100_190802_FIXED_rere.mzML | 16340 | HCD | 607.832 | 0 | 0.20 | 2 | K | EK+226.078ALVSSVR | Q | sp\|P07724\|ALBU_MOUSE | 98 | 95 | 1.34E-11 | 3.14E-04 |
| Issac_4_1_NAQ444_J100_190802_FIXED_rere.mzML | 16565 | HCD | 788.635 | 0 | 0.23 | 4 | R | VCLLHEKTPVSEHVTK+226.078CCSGSLVER | R | sp\|P07724\|ALBU_MOUSE | 190 | 156 | 2.47E-22 | 6.36E-15 |
| Issac_4_1_NAQ444_J100_190802_FIXED_rere.mzML | 16567 | HCD | 631.109 | 0 | 0.19 | 5 | R | VCLLHEK+226.078TPVSEHVTKCCSGSLVER | R | sp\|P07724\|ALBU_MOUSE | 160 | 65 | 2.09E-13 | 5.40E-06 |
| Issac_4_1_NAQ444_J100_190802_FIXED_rere.mzML | 16607 | HCD | 607.832 | 0 | 0.40 | 2 | K | EK+226.078ALVSSVR | Q | sp\|P07724\|ALBU_MOUSE | 85 | 83 | 1.34E-11 | 3.14E-04 |
| Issac_4_1_NAQ444_J100_190802_FIXED_rere.mzML | 16670 | HCD | 861.913 | 0 | -0.21 | 2 | K | NLVK+226.078TNCDLYEK | L | sp\|P07724\|ALBU_MOUSE | 157 | 146 | 7.04E-16 | 1.69E-08 |
| Issac_4_1_NAQ444_J100_190802_FIXED_rere.mzML | 16674 | HCD | 574.944 | 0 | -0.32 | 3 | K | NLVK+226.078TNCDLYEK | L | sp\|P07724\|ALBU_MOUSE | 147 | 143 | 4.04E-16 | 9.70E-09 |
| Issac_4_1_NAQ444_J100_190802_FIXED_rere.mzML | 16875 | HCD | 526.524 | 0 | 1.04 | 4 | R | VCLLHEK+226.078TPVSEHVTK | C | sp\|P07724\|ALBU_MOUSE | 176 | 162 | 1.31E-20 | 3.23E-13 |
| Issac_4_1_NAQ444_J100_190802_FIXED_rere.mzML | 16902 | HCD | 701.696 | 0 | 0.61 | 3 | R | VCLLHEK+226.078TPVSEHVTK | C | sp\|P07724\|ALBU_MOUSE | 162 | 95 | 7.55E-13 | 1.86E-05 |
| Issac_4_1_NAQ444_J100_190802_FIXED_rere.mzML | 16922 | HCD | 861.914 | 0 | 1.06 | 2 | K | NLVK+226.078TNCDLYEK | L | sp\|P07724\|ALBU_MOUSE | 136 | 134 | 2.79E-16 | 6.69E-09 |
| Issac_4_1_NAQ444_J100_190802_FIXED_rere.mzML | 18669 | HCD | 576.324 | 0 | 2.22 | 3 | K | LDGVKEK+226.078ALVSSVR | Q | sp\|P07724\|ALBU_MOUSE | 180 | 170 | 1.05E-17 | 2.55E-10 |
| Issac_4_1_NAQ444_J100_190802_FIXED_rere.mzML | 18748 | HCD | 863.981 | 0 | 0.99 | 2 | K | LDGVKEK+226.078ALVSSVR | Q | sp\|P07724\|ALBU_MOUSE | 66 | 46 | 3.33E-11 | 8.11E-04 |
| Issac_4_1_NAQ444_J100_190802_FIXED_rere.mzML | 18784 | HCD | 790.376 | 0 | 0.77 | 4 | K | YM(ox)CENQATISSK+226.078LQTCCDKPLLKK | A | sp\|P07724\|ALBU_MOUSE | 222 | 196 | 1.73E-24 | 4.44E-17 |
| Issac_4_1_NAQ444_J100_190802_FIXED_rere.mzML | 18819 | HCD | 1053.499 | 0 | 0.35 | 3 | K | YM(ox)CENQATISSK+226.078LQTCCDKPLLKK | A | sp\|P07724\|ALBU_MOUSE | 264 | 238 | 2.02E-25 | 5.18E-18 |
| Issac_4_1_NAQ444_J100_190802_FIXED_rere.mzML | 18952 | HCD | 576.323 | 0 | 0.95 | 3 | K | LDGVKEK+226.078ALVSSVR | Q | sp\|P07724\|ALBU_MOUSE | 117 | 87 | 1.11E-13 | 2.71E-06 |
| Issac_4_1_NAQ444_J100_190802_FIXED_rere.mzML | 19083 | HCD | 790.376 | 0 | -0.08 | 4 | K | YM(ox)CENQATISSK+226.078LQTCCDKPLLKK | A | sp\|P07724\|ALBU_MOUSE | 231 | 170 | 2.22E-19 | 5.70E-12 |
| Issac_4_1_NAQ444_J100_190802_FIXED_rere.mzML | 19110 | HCD | 1010.800 | 0 | -0.30 | 3 | K | YM(ox)CENQATISSK+226.078LQTCCDKPLLK | K | sp\|P07724\|ALBU_MOUSE | 255 | 84 | 5.88E-11 | 1.50E-03 |
| Issac_4_1_NAQ444_J100_190802_FIXED_rere.mzML | 19189 | HCD | 534.599 | 0 | -1.14 | 3 | K | LQTCCDK+226.078PLLK | K | sp\|P07724\|ALBU_MOUSE | 109 | 93 | 9.55E-13 | 2.27E-05 |
| Issac_4_1_NAQ444_J100_190802_FIXED_rere.mzML | 19194 | HCD | 801.396 | 0 | 0.38 | 2 | K | LQTCCDK+226.078PLLK | K | sp\|P07724\|ALBU_MOUSE | 143 | 136 | 1.95E-14 | 4.64E-07 |
| Issac_4_1_NAQ444_J100_190802_FIXED_rere.mzML | 19451 | HCD | 801.397 | 0 | 1.29 | 2 | K | LQTCCDK+226.078PLLK | K | sp\|P07724\|ALBU_MOUSE | 123 | 118 | 4.74E-14 | 1.13E-06 |
| Issac_4_1_NAQ444_J100_190802_FIXED_rere.mzML | 19525 | HCD | 790.376 | 0 | -0.23 | 4 | K | YM(ox)CENQATISSK+226.078LQTCCDKPLLKK | A | sp\|P07724\|ALBU_MOUSE | 126 | 68 | 2.74E-12 | 7.04E-05 |
| Issac_4_1_NAQ444_J100_190802_FIXED_rere.mzML | 19818 | HCD | 790.376 | 0 | 0.23 | 4 | K | YM(ox)CENQATISSK+226.078LQTCCDKPLLKK | A | sp\|P07724\|ALBU_MOUSE | 174 | 121 | 1.49E-17 | 3.82E-10 |
| Issac_4_1_NAQ444_J100_190802_FIXED_rere.mzML | 19846 | HCD | 786.627 | 1 | -0.76 | 4 | K | YMCENQATISSK+226.078LQTCCDKPLLKK | A | sp\|P07724\|ALBU_MOUSE | 126 | 66 | 1.68E-12 | 4.31E-05 |
| Issac_4_1_NAQ444_J100_190802_FIXED_rere.mzML | 19983 | HCD | 790.626 | 1 | -0.44 | 4 | K | YM(ox)CENQATISSK+226.078LQTCCDKPLLKK | A | sp\|P07724\|ALBU_MOUSE | 128 | 83 | 2.55E-14 | 6.56E-07 |
| Issac_4_1_NAQ444_J100_190802_FIXED_rere.mzML | 20298 | HCD | 729.380 | 0 | -1.26 | 3 | R | YTQK+226.078APQVSTPTLVEAAR | N | sp\|P07724\|ALBU_MOUSE | 154 | 145 | 3.60E-21 | 8.97E-14 |
| Issac_4_1_NAQ444_J100_190802_FIXED_rere.mzML | 20315 | HCD | 1093.568 | 0 | 0.67 | 2 | R | YTQK+226.078APQVSTPTLVEAAR | N | sp\|P07724\|ALBU_MOUSE | 193 | 173 | 6.34E-20 | 1.58E-12 |
| Issac_4_1_NAQ444_J100_190802_FIXED_rere.mzML | 20566 | HCD | 729.381 | 0 | -0.17 | 3 | R | YTQK+226.078APQVSTPTLVEAAR | N | sp\|P07724\|ALBU_MOUSE | 158 | 147 | 1.08E-20 | 2.70E-13 |
| Issac_4_1_NAQ444_J100_190802_FIXED_rere.mzML | 20579 | HCD | 1093.567 | 0 | -0.22 | 2 | R | YTQK+226.078APQVSTPTLVEAAR | N | sp\|P07724\|ALBU_MOUSE | 100 | 78 | 4.56E-15 | 1.14E-07 |
| Issac_4_1_NAQ444_J100_190802_FIXED_rere.mzML | 20954 | HCD | 729.381 | 0 | -0.17 | 3 | R | YTQK+226.078APQVSTPTLVEAAR | N | sp\|P07724\|ALBU_MOUSE | 127 | 66 | 1.75E-12 | 4.36E-05 |
| Issac_4_1_NAQ444_J100_190802_FIXED_rere.mzML | 21041 | HCD | 1011.137 | 1 | 2.01 | 3 | K | YM(ox)CENQATISSK+226.078LQTCCDKPLLK | K | sp\|P07724\|ALBU_MOUSE | 311 | 310 | 8.46E-30 | 2.16E-22 |
| Issac_4_1_NAQ444_J100_190802_FIXED_rere.mzML | 21050 | HCD | 1010.801 | 0 | 0.66 | 3 | K | YM(ox)CENQATISSK+226.078LQTCCDKPLLK | K | sp\|P07724\|ALBU_MOUSE | 306 | 305 | 8.46E-30 | 2.16E-22 |
| Issac_4_1_NAQ444_J100_190802_FIXED_rere.mzML | 21073 | HCD | 758.354 | 0 | 2.09 | 4 | K | YM(ox)CENQATISSK+226.078LQTCCDKPLLK | K | sp\|P07724\|ALBU_MOUSE | 258 | 187 | 6.75E-21 | 1.72E-13 |
| Issac_4_1_NAQ444_J100_190802_FIXED_rere.mzML | 21302 | HCD | 1010.801 | 0 | 0.42 | 3 | K | YM(ox)CENQATISSK+226.078LQTCCDKPLLK | K | sp\|P07724\|ALBU_MOUSE | 291 | 279 | 6.76E-28 | 1.73E-20 |
| Issac_4_1_NAQ444_J100_190802_FIXED_rere.mzML | 21326 | HCD | 758.352 | 0 | 0.08 | 4 | K | YM(ox)CENQATISSK+226.078LQTCCDKPLLK | K | sp\|P07724\|ALBU_MOUSE | 253 | 181 | 4.03E-20 | 1.03E-12 |
| Issac_4_1_NAQ444_J100_190802_FIXED_rere.mzML | 21521 | HCD | 758.603 | 1 | -0.14 | 4 | K | YM(ox)CENQATISSK+226.078LQTCCDKPLLK | K | sp\|P07724\|ALBU_MOUSE | 125 | 49 | 8.87E-13 | 2.27E-05 |
| Issac_4_1_NAQ444_J100_190802_FIXED_rere.mzML | 21561 | HCD | 1010.799 | 0 | -0.78 | 3 | K | YM(ox)CENQATISSK+226.078LQTCCDKPLLK | K | sp\|P07724\|ALBU_MOUSE | 160 | 128 | 5.28E-21 | 1.35E-13 |
| Issac_4_1_NAQ444_J100_190802_FIXED_rere.mzML | 21908 | HCD | 1011.135 | 1 | 0.14 | 3 | K | YM(ox)CENQATISSK+226.078LQTCCDKPLLK | K | sp\|P07724\|ALBU_MOUSE | 94 | 53 | 1.77E-16 | 4.53E-09 |
| Issac_4_1_NAQ444_J100_190802_FIXED_rere.mzML | 22031 | HCD | 933.423 | 1 | 1.59 | 3 | K | LVQEVTDFAK+226.078TCVADESAANCDK | S | sp\|P07724\|ALBU_MOUSE | 225 | 220 | 1.89E-28 | 4.83E-21 |
| Issac_4_1_NAQ444_J100_190802_FIXED_rere.mzML | 22038 | HCD | 933.088 | 0 | 0.46 | 3 | K | LVQEVTDFAK+226.078TCVADESAANCDK | S | sp\|P07724\|ALBU_MOUSE | 206 | 198 | 1.31E-26 | 3.34E-19 |
| Issac_4_1_NAQ444_J100_190802_FIXED_rere.mzML | 22093 | HCD | 1005.803 | 1 | 0.39 | 3 | K | YMCENQATISSK+226.078LQTCCDKPLLK | K | sp\|P07724\|ALBU_MOUSE | 110 | 66 | 4.21E-16 | 1.08E-08 |
| Issac_4_1_NAQ444_J100_190802_FIXED_rere.mzML | 22105 | HCD | 889.060 | 0 | -1.10 | 3 | K | TCVADESAANCDK+226.078SLHTLFGDK | L | sp\|P07724\|ALBU_MOUSE | 272 | 269 | 6.58E-28 | 1.67E-20 |
| Issac_4_1_NAQ444_J100_190802_FIXED_rere.mzML | 22141 | HCD | 1010.801 | 0 | 1.15 | 3 | K | YM(ox)CENQATISSK+226.078LQTCCDKPLLK | K | sp\|P07724\|ALBU_MOUSE | 211 | 62 | 1.33E-11 | 3.40E-04 |
| Issac_4_1_NAQ444_J100_190802_FIXED_rere.mzML | 22143 | HCD | 1010.801 | 0 | 1.15 | 3 | K | YM(ox)CENQATISSK+226.078LQTCCDKPLLK | K | sp\|P07724\|ALBU_MOUSE | 154 | 91 | 3.36E-17 | 8.58E-10 |
| Issac_4_1_NAQ444_J100_190802_FIXED_rere.mzML | 22162 | HCD | 1005.468 | 0 | -0.73 | 3 | K | YMCENQATISSK+226.078LQTCCDKPLLK | K | sp\|P07724\|ALBU_MOUSE | 154 | 114 | 7.83E-18 | 2.00E-10 |
| Issac_4_1_NAQ444_J100_190802_FIXED_rere.mzML | 22292 | HCD | 933.088 | 0 | 0.59 | 3 | K | LVQEVTDFAK+226.078TCVADESAANCDK | S | sp\|P07724\|ALBU_MOUSE | 211 | 205 | 4.53E-27 | 1.16E-19 |
| Issac_4_1_NAQ444_J100_190802_FIXED_rere.mzML | 22501 | HCD | 623.332 | 0 | -0.29 | 2 | R | AFK+226.078AWAVAR | L | sp\|P07724\|ALBU_MOUSE | 130 | 112 | 4.33E-13 | 1.01E-05 |
| Issac_4_1_NAQ444_J100_190802_FIXED_rere.mzML | 22603 | HCD | 933.088 | 0 | 0.52 | 3 | K | LVQEVTDFAK+226.078TCVADESAANCDK | S | sp\|P07724\|ALBU_MOUSE | 175 | 158 | 8.06E-25 | 2.06E-17 |
| Issac_4_1_NAQ444_J100_190802_FIXED_rere.mzML | 22669 | HCD | 933.422 | 1 | 0.02 | 3 | K | LVQEVTDFAK+226.078TCVADESAANCDK | S | sp\|P07724\|ALBU_MOUSE | 141 | 136 | 1.09E-24 | 2.79E-17 |
| Issac_4_1_NAQ444_J100_190802_FIXED_rere.mzML | 22751 | HCD | 623.332 | 0 | 0.78 | 2 | R | AFK+226.078AWAVAR | L | sp\|P07724\|ALBU_MOUSE | 119 | 97 | 1.90E-12 | 4.43E-05 |
| Issac_4_1_NAQ444_J100_190802_FIXED_rere.mzML | 22752 | HCD | 663.877 | 0 | 0.74 | 2 | K | K+226.078QTALAELVK | H | sp\|P07724\|ALBU_MOUSE | 121 | 121 | 3.95E-13 | 9.32E-06 |
| Issac_4_1_NAQ444_J100_190802_FIXED_rere.mzML | 23005 | HCD | 623.332 | 0 | 0.10 | 2 | R | AFK+226.078AWAVAR | L | sp\|P07724\|ALBU_MOUSE | 98 | 86 | 1.51E-12 | 3.52E-05 |
| Issac_4_1_NAQ444_J100_190802_FIXED_rere.mzML | 23257 | HCD | 623.332 | 0 | 0.49 | 2 | R | AFK+226.078AWAVAR | L | sp\|P07724\|ALBU_MOUSE | 94 | 91 | 1.17E-12 | 2.73E-05 |
| Issac_4_1_NAQ444_J100_190802_FIXED_rere.mzML | 23503 | HCD | 623.331 | 0 | -1.18 | 2 | R | AFK+226.078AWAVAR | L | sp\|P07724\|ALBU_MOUSE | 104 | 95 | 1.72E-12 | 4.01E-05 |
| Issac_4_1_NAQ444_J100_190802_FIXED_rere.mzML | 23758 | HCD | 623.332 | 0 | 0.39 | 2 | R | AFK+226.078AWAVAR | L | sp\|P07724\|ALBU_MOUSE | 101 | 96 | 1.71E-12 | 4.00E-05 |
| Issac_4_1_NAQ444_J100_190802_FIXED_rere.mzML | 24015 | HCD | 623.332 | 0 | 0.39 | 2 | R | AFK+226.078AWAVAR | L | sp\|P07724\|ALBU_MOUSE | 101 | 99 | 1.55E-12 | 3.62E-05 |
| Issac_4_1_NAQ444_J100_190802_FIXED_rere.mzML | 24317 | HCD | 623.332 | 0 | -0.20 | 2 | R | AFK+226.078AWAVAR | L | sp\|P07724\|ALBU_MOUSE | 96 | 94 | 1.45E-12 | 3.40E-05 |
| Issac_4_1_NAQ444_J100_190802_FIXED_rere.mzML | 24591 | HCD | 623.331 | 0 | -1.47 | 2 | R | AFK+226.078AWAVAR | L | sp\|P07724\|ALBU_MOUSE | 92 | 89 | 1.45E-12 | 3.40E-05 |
| Issac_4_1_NAQ444_J100_190802_FIXED_rere.mzML | 24879 | HCD | 623.331 | 0 | -1.96 | 2 | R | AFK+226.078AWAVAR | L | sp\|P07724\|ALBU_MOUSE | 95 | 88 | 1.17E-12 | 2.72E-05 |
| Issac_4_1_NAQ444_J100_190802_FIXED_rere.mzML | 25195 | HCD | 789.364 | 0 | -1.16 | 3 | K | CSYDEHAK+226.078LVQEVTDFAK | T | sp\|P07724\|ALBU_MOUSE | 207 | 160 | 1.93E-20 | 4.80E-13 |
| Issac_4_1_NAQ444_J100_190802_FIXED_rere.mzML | 25430 | HCD | 623.332 | 0 | -0.29 | 2 | R | AFK+226.078AWAVAR | L | sp\|P07724\|ALBU_MOUSE | 87 | 82 | 2.29E-12 | 5.34E-05 |
| Issac_4_1_NAQ444_J100_190802_FIXED_rere.mzML | 25744 | HCD | 623.332 | 0 | 0.20 | 2 | R | AFK+226.078AWAVAR | L | sp\|P07724\|ALBU_MOUSE | 82 | 79 | 1.46E-12 | 3.40E-05 |
| Issac_4_1_NAQ444_J100_190802_FIXED_rere.mzML | 26020 | HCD | 623.332 | 0 | 0.59 | 2 | R | AFK+226.078AWAVAR | L | sp\|P07724\|ALBU_MOUSE | 94 | 91 | 1.45E-12 | 3.40E-05 |
| Issac_4_1_NAQ444_J100_190802_FIXED_rere.mzML | 26098 | HCD | 838.746 | 1 | -0.19 | 3 | R | RPCFSALTVDETYVPK+226.078EFK | A | sp\|P07724\|ALBU_MOUSE | 83 | 43 | 7.12E-13 | 1.79E-05 |
| Issac_4_1_NAQ444_J100_190802_FIXED_rere.mzML | 26115 | HCD | 838.413 | 0 | 2.11 | 3 | R | RPCFSALTVDETYVPK+226.078EFK | A | sp\|P07724\|ALBU_MOUSE | 95 | 45 | 2.79E-11 | 7.00E-04 |
| Issac_4_1_NAQ444_J100_190802_FIXED_rere.mzML | 26536 | HCD | 623.332 | 0 | 0.20 | 2 | R | AFK+226.078AWAVAR | L | sp\|P07724\|ALBU_MOUSE | 95 | 91 | 1.45E-12 | 3.40E-05 |
| Issac_4_1_NAQ444_J100_190802_FIXED_rere.mzML | 26918 | HCD | 623.331 | 0 | -0.88 | 2 | R | AFK+226.078AWAVAR | L | sp\|P07724\|ALBU_MOUSE | 82 | 79 | 1.45E-12 | 3.40E-05 |
| Issac_4_1_NAQ444_J100_190802_FIXED_rere.mzML | 27920 | HCD | 623.332 | 0 | -0.29 | 2 | R | AFK+226.078AWAVAR | L | sp\|P07724\|ALBU_MOUSE | 101 | 97 | 1.02E-12 | 2.37E-05 |
| Issac_4_1_NAQ444_J100_190802_FIXED_rere.mzML | 28100 | HCD | 901.428 | 0 | 1.56 | 4 | K | TCVADESAANCDK+226.078SLHTLFGDKLCAIPNLR | E | sp\|P07724\|ALBU_MOUSE | 313 | 302 | 1.29E-35 | 3.40E-28 |
| Issac_4_1_NAQ444_J100_190802_FIXED_rere.mzML | 28151 | HCD | 1201.570 | 0 | 2.34 | 3 | K | TCVADESAANCDK+226.078SLHTLFGDKLCAIPNLR | E | sp\|P07724\|ALBU_MOUSE | 341 | 316 | 3.72E-33 | 9.79E-26 |
| Issac_4_1_NAQ444_J100_190802_FIXED_rere.mzML | 28363 | HCD | 901.427 | 0 | -0.27 | 4 | K | TCVADESAANCDK+226.078SLHTLFGDKLCAIPNLR | E | sp\|P07724\|ALBU_MOUSE | 331 | 304 | 3.81E-33 | 1.00E-25 |
| Issac_4_1_NAQ444_J100_190802_FIXED_rere.mzML | 28419 | HCD | 1014.696 | 0 | 0.48 | 4 | K | TVM(ox)DDFAQFLDTCCK+226.078AADKDTCFSTEGPNLVTR | C | sp\|P07724\|ALBU_MOUSE | 128 | 74 | 6.01E-17 | 1.60E-09 |
| Issac_4_1_NAQ444_J100_190802_FIXED_rere.mzML | 28462 | HCD | 901.675 | 1 | -3.57 | 4 | K | TCVADESAANCDK+226.078SLHTLFGDKLCAIPNLR | E | sp\|P07724\|ALBU_MOUSE | 301 | 282 | 1.41E-33 | 3.72E-26 |
| Issac_4_1_NAQ444_J100_190802_FIXED_rere.mzML | 30358 | HCD | 901.677 | 1 | -0.52 | 4 | K | TCVADESAANCDKSLHTLFGDK+226.078LCAIPNLR | E | sp\|P07724\|ALBU_MOUSE | 226 | 183 | 5.91E-23 | 1.56E-15 |
| Issac_4_1_NAQ444_J100_190802_FIXED_rere.mzML | 30402 | HCD | 901.428 | 0 | 0.68 | 4 | K | TCVADESAANCDKSLHTLFGDK+226.078LCAIPNLR | E | sp\|P07724\|ALBU_MOUSE | 250 | 170 | 2.57E-20 | 6.75E-13 |
| Issac_4_1_NAQ444_J100_190802_FIXED_rere.mzML | 30451 | HCD | 906.428 | 1 | -2.54 | 4 | K | SLHTLFGDK+226.078LCAIPNLRENYGELADCCTK | Q | sp\|P07724\|ALBU_MOUSE | 208 | 119 | 4.51E-16 | 1.18E-08 |
| Issac_4_1_NAQ444_J100_190802_FIXED_rere.mzML | 30680 | HCD | 1061.870 | 0 | 0.00 | 3 | K | NLVK+226.078TNCDLYEKLGEYGFQNAILVR | Y | sp\|P07724\|ALBU_MOUSE | 323 | 318 | 2.75E-31 | 7.10E-24 |
| Issac_4_1_NAQ444_J100_190802_FIXED_rere.mzML | 30713 | HCD | 796.655 | 0 | 0.08 | 4 | K | NLVK+226.078TNCDLYEKLGEYGFQNAILVR | Y | sp\|P07724\|ALBU_MOUSE | 187 | 110 | 1.56E-16 | 4.01E-09 |
| Issac_4_1_NAQ444_J100_190802_FIXED_rere.mzML | 30800 | HCD | 1062.204 | 1 | -0.78 | 3 | K | NLVK+226.078TNCDLYEKLGEYGFQNAILVR | Y | sp\|P07724\|ALBU_MOUSE | 255 | 232 | 9.07E-27 | 2.34E-19 |
| Issac_4_1_NAQ444_J100_190802_FIXED_rere.mzML | 31650 | HCD | 727.716 | 0 | 1.01 | 3 | K | SLHTLFGDK+226.078LCAIPNLR | E | sp\|P07724\|ALBU_MOUSE | 162 | 108 | 8.33E-14 | 2.07E-06 |
| Issac_4_1_NAQ444_J100_190802_FIXED_rere.mzML | 31688 | HCD | 1091.068 | 0 | -1.12 | 2 | K | SLHTLFGDK+226.078LCAIPNLR | E | sp\|P07724\|ALBU_MOUSE | 187 | 161 | 5.47E-18 | 1.36E-10 |
| Issac_4_1_NAQ444_J100_190802_FIXED_rere.mzML | 31904 | HCD | 727.715 | 0 | 0.25 | 3 | K | SLHTLFGDK+226.078LCAIPNLR | E | sp\|P07724\|ALBU_MOUSE | 159 | 102 | 1.81E-13 | 4.48E-06 |
| Issac_4_1_NAQ444_J100_190802_FIXED_rere.mzML | 32287 | HCD | 945.423 | 0 | 1.10 | 3 | K | ATAEQLK+226.078TVM(ox)DDFAQFLDTCCK | A | sp\|P07724\|ALBU_MOUSE | 159 | 127 | 3.69E-18 | 9.38E-11 |
| Issac_4_1_NAQ444_J100_190802_FIXED_rere.mzML | 32610 | HCD | 727.714 | 0 | -1.17 | 3 | K | SLHTLFGDK+226.078LCAIPNLR | E | sp\|P07724\|ALBU_MOUSE | 106 | 47 | 7.11E-11 | 1.76E-03 |
| Issac_4_1_NAQ444_J100_190802_FIXED_rere.mzML | 32869 | HCD | 910.441 | 0 | 0.87 | 3 | K | TNCDLYEK+226.078LGEYGFQNAILVR | Y | sp\|P07724\|ALBU_MOUSE | 211 | 200 | 1.78E-24 | 4.51E-17 |
| Issac_4_1_NAQ444_J100_190802_FIXED_rere.mzML | 33142 | HCD | 910.442 | 0 | 2.01 | 3 | K | TNCDLYEK+226.078LGEYGFQNAILVR | Y | sp\|P07724\|ALBU_MOUSE | 105 | 53 | 1.78E-13 | 4.51E-06 |
| Issac_4_1_NAQ444_J100_190802_FIXED_rere.mzML | 34836 | HCD | 1145.552 | 1 | -0.19 | 3 | R | VGTK+226.078CCTLPEDQRLPCVEDYLSAILNR | V | sp\|P07724\|ALBU_MOUSE | 267 | 228 | 3.08E-26 | 8.02E-19 |
| Issac_4_1_NAQ444_J100_190802_FIXED_rere.mzML | 34872 | HCD | 1145.217 | 0 | -0.53 | 3 | R | VGTK+226.078CCTLPEDQRLPCVEDYLSAILNR | V | sp\|P07724\|ALBU_MOUSE | 271 | 253 | 5.15E-30 | 1.34E-22 |
| Issac_4_1_NAQ444_J100_190802_FIXED_rere.mzML | 34873 | HCD | 859.164 | 0 | -0.50 | 4 | R | VGTK+226.078CCTLPEDQRLPCVEDYLSAILNR | V | sp\|P07724\|ALBU_MOUSE | 252 | 188 | 2.07E-22 | 5.37E-15 |
| Issac_4_1_NAQ444_J100_190802_FIXED_rere.mzML | 35211 | HCD | 1211.242 | 1 | -0.18 | 3 | K | CCAEANPPACYGTVLAEFQPLVEEPK+226.078NLVK | T | sp\|P07724\|ALBU_MOUSE | 193 | 101 | 1.67E-16 | 4.38E-09 |
| Issac_4_1_NAQ444_J100_190802_FIXED_rere.mzML | 35244 | HCD | 899.445 | 0 | -0.54 | 3 | K | GLVLIAFSQYLQK+226.078CSYDEHAK | L | sp\|P07724\|ALBU_MOUSE | 217 | 205 | 2.32E-24 | 5.87E-17 |
| Issac_4_1_NAQ444_J100_190802_FIXED_rere.mzML | 35263 | HCD | 1210.907 | 0 | -0.71 | 3 | K | CCAEANPPACYGTVLAEFQPLVEEPK+226.078NLVK | T | sp\|P07724\|ALBU_MOUSE | 181 | 85 | 4.96E-15 | 1.31E-07 |
| Issac_4_1_NAQ444_J100_190802_FIXED_rere.mzML | 37828 | HCD | 838.065 | 0 | -1.46 | 3 | K | NYAEAK+226.078DVFLGTFLYEYSR | R | sp\|P07724\|ALBU_MOUSE | 126 | 77 | 1.95E-14 | 4.89E-07 |
| Issac_4_1_NAQ444_J100_190802_FIXED_rere.mzML | 38961 | HCD | 817.750 | 1 | 1.74 | 3 | S | QTFPNADFAEITK+226.078LATDLTK | V | sp\|P07724\|ALBU_MOUSE | 97 | 52 | 8.89E-12 | 2.24E-04 |
| Issac_4_1_NAQ444_J100_190802_FIXED_rere.mzML | 39066 | HCD | 884.122 | 0 | 2.90 | 3 | R | LSQTFPNADFAEITK+226.078LATDLTK | V | sp\|P07724\|ALBU_MOUSE | 102 | 33 | 6.49E-11 | 1.65E-03 |
| Issac_4_1_NAQ444_J100_190802_FIXED_rere.mzML | 39140 | HCD | 997.854 | 0 | -0.98 | 3 | R | LSQTFPNADFAEITK+226.078LATDLTKVNK | E | sp\|P07724\|ALBU_MOUSE | 174 | 119 | 6.73E-17 | 1.74E-09 |
| Issac_4_1_NAQ444_J100_190802_FIXED_rere.mzML | 39870 | HCD | 884.119 | 0 | -0.35 | 3 | R | LSQTFPNADFAEITK+226.078LATDLTK | V | sp\|P07724\|ALBU_MOUSE | 240 | 214 | 1.09E-23 | 2.78E-16 |
| Issac_4_1_NAQ444_J100_190802_FIXED_rere.mzML | 39892 | HCD | 1325.675 | 0 | -0.46 | 2 | R | LSQTFPNADFAEITK+226.078LATDLTK | V | sp\|P07724\|ALBU_MOUSE | 185 | 153 | 4.51E-20 | 1.15E-12 |
| Issac_4_1_NAQ444_J100_190802_FIXED_rere.mzML | 40122 | HCD | 884.120 | 0 | 0.35 | 3 | R | LSQTFPNADFAEITK+226.078LATDLTK | V | sp\|P07724\|ALBU_MOUSE | 234 | 223 | 2.60E-26 | 6.61E-19 |
| Issac_4_1_NAQ444_J100_190802_FIXED_rere.mzML | 40145 | HCD | 1325.676 | 0 | 0.28 | 2 | R | LSQTFPNADFAEITK+226.078LATDLTK | V | sp\|P07724\|ALBU_MOUSE | 218 | 192 | 4.07E-22 | 1.04E-14 |
| Issac_4_1_NAQ444_J100_190802_FIXED_rere.mzML | 40333 | HCD | 979.838 | 0 | 0.44 | 3 | R | YNDLGEQHFK+226.078GLVLIAFSQYLQK | C | sp\|P07724\|ALBU_MOUSE | 214 | 135 | 1.21E-15 | 3.10E-08 |
| Issac_4_1_NAQ444_J100_190802_FIXED_rere.mzML | 40375 | HCD | 884.119 | 0 | -0.21 | 3 | R | LSQTFPNADFAEITK+226.078LATDLTK | V | sp\|P07724\|ALBU_MOUSE | 244 | 213 | 2.56E-22 | 6.52E-15 |

| | | | | | | | | | | | | | | | |
|---|---|---|---|---|---|---|---|---|---|---|---|---|---|---|---|
| Issac_4_1_NAQ444_J100_190802_FIXED_rere.mzML | 40544 | HCD | | 980.171 | 1 | -0.73 | 3 | R | YNDLGEQHFK+226.078GLVLIAFSQYLQK | C | sp\|P07724\|ALBU_MOUSE | 118 | 76 | 1.92E-13 | 4.92E-06 |
| Issac_4_1_NAQ444_J100_190802_FIXED_rere.mzML | 40629 | HCD | | 884.118 | 0 | -1.10 | 3 | R | LSQTFPNADFAEITK+226.078LATDLTK | V | sp\|P07724\|ALBU_MOUSE | 188 | 165 | 3.17E-19 | 8.06E-12 |
| Issac_4_1_NAQ444_J100_190802_FIXED_rere.mzML | 40934 | HCD | | 884.116 | 0 | -4.07 | 3 | R | LSQTFPNADFAEITK+226.078LATDLTK | V | sp\|P07724\|ALBU_MOUSE | 112 | 73 | 8.33E-15 | 2.12E-07 |
| Issac_4_1_NAQ444_J100_190802_FIXED_rere.mzML | 40955 | HCD | | 884.452 | 1 | -1.77 | 3 | R | LSQTFPNADFAEITK+226.078LATDLTK | V | sp\|P07724\|ALBU_MOUSE | 114 | 68 | 1.26E-13 | 3.20E-06 |
| Issac_4_1_NAQ444_J100_190802_FIXED_rere.mzML | 41511 | HCD | | 884.452 | 1 | -2.12 | 3 | R | LSQTFPNADFAEITK+226.078LATDLTK | V | sp\|P07724\|ALBU_MOUSE | 89 | 33 | 8.16E-11 | 2.08E-03 |
| Issac_4_1_NAQ444_J100_190802_FIXED_rere.mzML | 51478 | HCD | | 623.332 | 0 | -0.10 | 2 | R | AFK+226.078AWAVAR | L | sp\|P07724\|ALBU_MOUSE | 104 | 101 | 1.45E-12 | 3.40E-05 |
| Issac_4_1_NAQ444_J100_190802_FIXED_rere.mzML | 51564 | HCD | | 1010.800 | 0 | -0.12 | 3 | K | YM(ox)CENQATISSK+226.078LQTCCDKPLLK | K | sp\|P07724\|ALBU_MOUSE | 124 | 55 | 2.56E-12 | 6.54E-05 |
| Issac_4_1_NAQ444_J100_190802_FIXED_rere.mzML | 51566 | HCD | | 1005.469 | 0 | 0.00 | 3 | K | YMCENQATISSK+226.078LQTCCDKPLLK | K | sp\|P07724\|ALBU_MOUSE | 160 | 93 | 5.35E-15 | 1.37E-07 |
| Issac_4_1_NAQ444_J100_190802_FIXED_rere.mzML | 51651 | HCD | | 790.375 | 0 | -0.93 | 4 | K | YM(ox)CENQATISSK+226.078LQTCCDKPLLKK | A | sp\|P07724\|ALBU_MOUSE | 135 | 75 | 1.46E-12 | 3.75E-05 |
| Issac_4_1_NAQ444_J100_190802_FIXED_rere.mzML | 17365 | HCD | | 716.862 | 0 | -0.60 | 2 | R | NVEK+226.078ISM(ox)LEK | I | sp\|P19221\|THRB_MOUSE | 108 | 103 | 1.48E-12 | 3.49E-05 |
| Issac_4_1_NAQ444_J100_190802_FIXED_rere.mzML | 20032 | HCD | | 576.638 | 0 | 1.16 | 3 | K | ISM(ox)LEK+226.078IYVHPR | Y | sp\|P19221\|THRB_MOUSE | 109 | 82 | 8.65E-14 | 2.08E-06 |
| Issac_4_1_NAQ444_J100_190802_FIXED_rere.mzML | 6719 | HCD | | 602.794 | 0 | -0.61 | 2 | R | TK+226.078SGVACQK | W | sp\|P20918\|PLMN_MOUSE | 84 | 71 | 1.28E-10 | 2.98E-03 |
| Issac_4_1_NAQ444_J100_190802_FIXED_rere.mzML | 19437 | HCD | | 779.373 | 0 | 0.31 | 3 | K | EQQCVIM(ox)AENSK+226.078TSSIIR | M | sp\|P20918\|PLMN_MOUSE | 108 | 97 | 9.47E-17 | 2.36E-09 |
| Issac_4_1_NAQ444_J100_190802_FIXED_rere.mzML | 13890 | HCD | | 711.019 | 0 | 0.43 | 3 | R | TK+226.078TPNTSPAELKDM(ox)VEK | H | sp\|P21614\|VTDB_MOUSE | 103 | 53 | 1.64E-11 | 4.06E-04 |
| Issac_4_1_NAQ444_J100_190802_FIXED_rere.mzML | 24449 | HCD | | 656.668 | 0 | 0.37 | 3 | R | K+226.078FSSSTFEQVNQLVK | E | sp\|P21614\|VTDB_MOUSE | 120 | 99 | 1.31E-15 | 3.21E-08 |
| Issac_4_1_NAQ444_J100_190802_FIXED_rere.mzML | 24458 | HCD | | 984.499 | 0 | 0.37 | 2 | R | K+226.078FSSSTFEQVNQLVK | E | sp\|P21614\|VTDB_MOUSE | 136 | 133 | 3.23E-18 | 7.92E-11 |
| Issac_4_1_NAQ444_J100_190802_FIXED_rere.mzML | 16108 | HCD | | 678.372 | 0 | 1.80 | 2 | K | GTQGK+226.078IVEAVK | E | sp\|P22599\|A1AT2_MOUSE | 120 | 116 | 5.34E-14 | 1.27E-06 |
| Issac_4_1_NAQ444_J100_190802_FIXED_rere.mzML | 21770 | HCD | | 767.416 | 0 | 0.72 | 3 | K | VINDFVEKGTQGK+226.078IVEAVK | E | sp\|P22599\|A1AT2_MOUSE | 149 | 122 | 1.83E-17 | 4.59E-10 |
| Issac_4_1_NAQ444_J100_190802_FIXED_rere.mzML | 29548 | HCD | | 712.918 | 0 | -0.34 | 2 | K | ELISK+226.078ILLNR | R | sp\|P22599\|A1AT2_MOUSE | 116 | 113 | 6.56E-12 | 1.55E-04 |
| Issac_4_1_NAQ444_J100_190802_FIXED_rere.mzML | 25520 | HCD | | 705.725 | 0 | -0.69 | 3 | G | HSLLPPVVDTTQGK+226.078VLGK | Y | sp\|P23953\|EST1C_MOUSE | 190 | 162 | 1.37E-18 | 3.41E-11 |
| Issac_4_1_NAQ444_J100_190802_FIXED_rere.mzML | 33450 | HCD | | 898.462 | 1 | -0.66 | 3 | K | APEEILAEK+226.078SFNTVPYIVGFNK | Q | sp\|P23953\|EST1C_MOUSE | 159 | 139 | 9.94E-23 | 2.53E-15 |
| Issac_4_1_NAQ444_J100_190802_FIXED_rere.mzML | 33493 | HCD | | 898.128 | 0 | 0.14 | 3 | K | APEEILAEK+226.078SFNTVPYIVGFNK | Q | sp\|P23953\|EST1C_MOUSE | 192 | 170 | 6.64E-24 | 1.69E-16 |
| Issac_4_1_NAQ444_J100_190802_FIXED_rere.mzML | 9515 | HCD | | 386.531 | 0 | -0.08 | 3 | R | ELNYK+226.078HK | D | sp\|P28665\|MUG1_MOUSE | 100 | 100 | 9.29E-11 | 2.11E-03 |
| Issac_4_1_NAQ444_J100_190802_FIXED_rere.mzML | 9542 | HCD | | 579.292 | 0 | 0.00 | 2 | R | ELNYK+226.078HK | D | sp\|P28665\|MUG1_MOUSE | 79 | 69 | 1.30E-10 | 2.94E-03 |
| Issac_4_1_NAQ444_J100_190802_FIXED_rere.mzML | 18861 | HCD | | 602.124 | 0 | 0.71 | 5 | K | NKESVVFVQTDKPVYKPGQSVK+226.078FR | V | sp\|P28665\|MUG1_MOUSE | 190 | 133 | 2.46E-18 | 6.31E-11 |
| Issac_4_1_NAQ444_J100_190802_FIXED_rere.mzML | 18880 | HCD | | 752.405 | 0 | 3.16 | 4 | K | NKESVVFVQTDKPVYKPGQSVK+226.078FR | V | sp\|P28665\|MUG1_MOUSE | 156 | 52 | 4.81E-11 | 1.23E-03 |
| Issac_4_1_NAQ444_J100_190802_FIXED_rere.mzML | 22321 | HCD | | 890.690 | 0 | 0.41 | 4 | K | VPNAM(ox)SVNDEVLSVTACGK+226.078YTYGKPVPGHVK | I | sp\|P28665\|MUG1_MOUSE | 214 | 123 | 6.80E-17 | 1.80E-09 |
| Issac_4_1_NAQ444_J100_190802_FIXED_rere.mzML | 24832 | HCD | | 566.321 | 0 | -0.22 | 2 | K | TK+226.078ALGFLR | A | sp\|P28665\|MUG1_MOUSE | 91 | 91 | 6.50E-11 | 1.50E-03 |
| Issac_4_1_NAQ444_J100_190802_FIXED_rere.mzML | 25924 | HCD | | 717.734 | 0 | 0.17 | 3 | R | LVDIKGDPIPNEK+226.078VFIK | A | sp\|P28665\|MUG1_MOUSE | 113 | 97 | 2.94E-16 | 7.30E-09 |
| Issac_4_1_NAQ444_J100_190802_FIXED_rere.mzML | 26924 | HCD | | 862.446 | 0 | -1.06 | 3 | K | QM(ox)SFSLAAEPIQGPYK+226.078IVVHK | E | sp\|P28665\|MUG1_MOUSE | 165 | 129 | 3.71E-19 | 9.38E-12 |
| Issac_4_1_NAQ444_J100_190802_FIXED_rere.mzML | 27874 | HCD | | 475.932 | 0 | 0.06 | 3 | R | HGIPFFVK+226.078VR | L | sp\|P28665\|MUG1_MOUSE | 78 | 68 | 1.70E-11 | 4.01E-04 |
| Issac_4_1_NAQ444_J100_190802_FIXED_rere.mzML | 29129 | HCD | | 983.812 | 0 | 1.61 | 3 | R | FNVDLK+226.078VPNAM(ox)SVNDEVLSVTACGK | Y | sp\|P28665\|MUG1_MOUSE | 154 | 92 | 6.46E-17 | 1.67E-09 |
| Issac_4_1_NAQ444_J100_190802_FIXED_rere.mzML | 29250 | HCD | | 658.394 | 0 | 0.09 | 2 | R | ITNK+226.078LIFLK | A | sp\|P28665\|MUG1_MOUSE | 122 | 109 | 2.12E-11 | 4.96E-04 |
| Issac_4_1_NAQ444_J100_190802_FIXED_rere.mzML | 29373 | HCD | | 658.896 | 1 | 0.33 | 2 | R | ITNK+226.078LIFLK | A | sp\|P28665\|MUG1_MOUSE | 122 | 101 | 2.62E-11 | 6.12E-04 |
| Issac_4_1_NAQ444_J100_190802_FIXED_rere.mzML | 33536 | HCD | | 1075.801 | 0 | 0.57 | 4 | R | AVDQSVLLLKPESELSPSWIYNLPGM(ox)QQNK+226.078FVPSSR | L | sp\|P28665\|MUG1_MOUSE | 168 | 86 | 1.54E-16 | 4.15E-09 |
| Issac_4_1_NAQ444_J100_190802_FIXED_rere.mzML | 33840 | HCD | | 793.422 | 0 | -0.15 | 4 | R | VVSM(ox)DK+226.078M(ox)LRPLNELLPLAYIEDPKK | N | sp\|P28665\|MUG1_MOUSE | 212 | 117 | 1.56E-14 | 4.02E-07 |
| Issac_4_1_NAQ444_J100_190802_FIXED_rere.mzML | 33922 | HCD | | 1057.560 | 0 | -0.12 | 3 | R | VVSM(ox)DK+226.078M(ox)LRPLNELLPLAYIEDPKK | N | sp\|P28665\|MUG1_MOUSE | 187 | 77 | 1.28E-12 | 3.30E-05 |
| Issac_4_1_NAQ444_J100_190802_FIXED_rere.mzML | 34111 | HCD | | 793.420 | 0 | -2.23 | 4 | R | VVSM(ox)DK+226.078M(ox)LRPLNELLPLAYIEDPKK | N | sp\|P28665\|MUG1_MOUSE | 213 | 119 | 1.72E-14 | 4.43E-07 |
| Issac_4_1_NAQ444_J100_190802_FIXED_rere.mzML | 36002 | HCD | | 1015.194 | 1 | -2.02 | 3 | R | VVSM(ox)DK+226.078M(ox)LRPLNELLPLAYIEDPK | K | sp\|P28665\|MUG1_MOUSE | 146 | 48 | 1.79E-11 | 4.59E-04 |
| Issac_4_1_NAQ444_J100_190802_FIXED_rere.mzML | 44753 | HCD | | 1012.531 | 0 | 1.27 | 3 | R | K+226.078NLFDELVLDKDLFQCVSFIIPR | L | sp\|P28665\|MUG1_MOUSE | 340 | 335 | 7.37E-29 | 1.88E-21 |
| Issac_4_1_NAQ444_J100_190802_FIXED_rere.mzML | 44868 | HCD | | 1012.862 | 1 | -1.43 | 3 | R | K+226.078NLFDELVLDKDLFQCVSFIIPR | L | sp\|P28665\|MUG1_MOUSE | 226 | 200 | 8.14E-23 | 2.08E-15 |
| Issac_4_1_NAQ444_J100_190802_FIXED_rere.mzML | 17567 | HCD | | 705.893 | 0 | 0.26 | 2 | K | QVLNQIDK+226.078VK | V | sp\|P29699\|FETUA_MOUSE | 148 | 148 | 3.93E-13 | 9.27E-06 |
| Issac_4_1_NAQ444_J100_190802_FIXED_rere.mzML | 21047 | HCD | | 812.402 | 0 | -0.30 | 2 | G | NPVDDICIAK+226.078PR | D | sp\|P32261\|ANT3_MOUSE | 182 | 158 | 3.56E-15 | 8.54E-08 |
| Issac_4_1_NAQ444_J100_190802_FIXED_rere.mzML | 30498 | HCD | | 704.006 | 0 | 1.04 | 3 | K | TSDQIHFFFAK+226.078LNCR | L | sp\|P32261\|ANT3_MOUSE | 178 | 157 | 2.52E-19 | 6.17E-12 |
| Issac_4_1_NAQ444_J100_190802_FIXED_rere.mzML | 30562 | HCD | | 704.339 | 1 | -0.92 | 3 | K | TSDQIHFFFAK+226.078LNCR | L | sp\|P32261\|ANT3_MOUSE | 160 | 142 | 5.93E-19 | 1.45E-11 |
| Issac_4_1_NAQ444_J100_190802_FIXED_rere.mzML | 14609 | HCD | | 537.267 | 0 | -0.23 | 3 | K | LYHAFSATK+226.078M(ox)AK | T | sp\|P97290\|IC1_MOUSE | 86 | 55 | 4.19E-12 | 1.01E-04 |
| Issac_4_1_NAQ444_J100_190802_FIXED_rere.mzML | 15549 | HCD | | 793.370 | 0 | 0.15 | 2 | Q | DEPQSQWDK+226.078VK | D | sp\|Q00623\|APOA1_MOUSE | 139 | 124 | 1.42E-13 | 3.38E-06 |
| Issac_4_1_NAQ444_J100_190802_FIXED_rere.mzML | 15584 | HCD | | 529.249 | 0 | -0.23 | 3 | Q | DEPQSQWDK+226.078VK | D | sp\|Q00623\|APOA1_MOUSE | 98 | 82 | 2.32E-12 | 5.52E-05 |
| Issac_4_1_NAQ444_J100_190802_FIXED_rere.mzML | 15807 | HCD | | 793.371 | 0 | 1.46 | 2 | Q | DEPQSQWDK+226.078VK | D | sp\|Q00623\|APOA1_MOUSE | 111 | 93 | 2.02E-12 | 4.81E-05 |
| Issac_4_1_NAQ444_J100_190802_FIXED_rere.mzML | 19074 | HCD | | 626.327 | 0 | 1.27 | 3 | K | QK+226.078VQPYLDEFQKK | W | sp\|Q00623\|APOA1_MOUSE | 116 | 94 | 4.14E-14 | 1.00E-06 |
| Issac_4_1_NAQ444_J100_190802_FIXED_rere.mzML | 23171 | HCD | | 703.687 | 0 | 0.87 | 3 | K | VK+226.078DFANVYVDAVKDSGR | D | sp\|Q00623\|APOA1_MOUSE | 135 | 101 | 3.93E-16 | 9.74E-09 |
| Issac_4_1_NAQ444_J100_190802_FIXED_rere.mzML | 24094 | HCD | | 847.435 | 0 | 0.22 | 2 | K | VK+226.078DFANVYVDAVK | D | sp\|Q00623\|APOA1_MOUSE | 161 | 157 | 5.05E-18 | 1.22E-10 |
| Issac_4_1_NAQ444_J100_190802_FIXED_rere.mzML | 29675 | HCD | | 936.450 | 0 | 0.46 | 3 | Q | DEPQSQWDKVK+226.078DFANVYVDAVK | D | sp\|Q00623\|APOA1_MOUSE | 235 | 207 | 6.67E-25 | 1.70E-17 |
| Issac_4_1_NAQ444_J100_190802_FIXED_rere.mzML | 29852 | HCD | | 936.783 | 1 | -0.82 | 3 | Q | DEPQSQWDK+226.078VKDFANVYVDAVK | D | sp\|Q00623\|APOA1_MOUSE | 210 | 191 | 1.62E-26 | 4.13E-19 |
| Issac_4_1_NAQ444_J100_190802_FIXED_rere.mzML | 29898 | HCD | | 702.589 | 0 | 0.00 | 4 | Q | DEPQSQWDK+226.078VKDFANVYVDAVK | D | sp\|Q00623\|APOA1_MOUSE | 208 | 109 | 1.54E-15 | 3.92E-08 |
| Issac_4_1_NAQ444_J100_190802_FIXED_rere.mzML | 29939 | HCD | | 936.449 | 0 | -0.26 | 3 | Q | DEPQSQWDK+226.078VKDFANVYVDAVK | D | sp\|Q00623\|APOA1_MOUSE | 195 | 169 | 1.98E-26 | 5.03E-19 |
| Issac_4_1_NAQ444_J100_190802_FIXED_rere.mzML | 30017 | HCD | | 806.385 | 0 | 0.08 | 4 | Q | DEPQSQWDK+226.078VKDFANVYVDAVKDSGR | D | sp\|Q00623\|APOA1_MOUSE | 210 | 171 | 2.38E-23 | 6.17E-16 |
| Issac_4_1_NAQ444_J100_190802_FIXED_rere.mzML | 30300 | HCD | | 936.449 | 0 | -0.72 | 3 | Q | DEPQSQWDKVK+226.078DFANVYVDAVK | D | sp\|Q00623\|APOA1_MOUSE | 114 | 72 | 2.84E-16 | 7.22E-09 |
| Issac_4_1_NAQ444_J100_190802_FIXED_rere.mzML | 30324 | HCD | | 936.784 | 1 | 0.15 | 3 | Q | DEPQSQWDKVK+226.078DFANVYVDAVK | D | sp\|Q00623\|APOA1_MOUSE | 163 | 132 | 2.37E-21 | 6.03E-14 |
| Issac_4_1_NAQ444_J100_190802_FIXED_rere.mzML | 41502 | HCD | | 818.115 | 0 | -0.30 | 3 | K | KLEQDTVFVLANYILFK+226.078GK | W | sp\|Q00898\|A1AT5_MOUSE | 159 | 114 | 2.47E-14 | 6.18E-07 |
| Issac_4_1_NAQ444_J100_190802_FIXED_rere.mzML | 32675 | HCD | | 932.127 | 0 | -0.46 | 3 | K | YCTAISGDLHILPVAFK+226.078GDSFTR | T | sp\|Q01279\|EGFR_MOUSE | 142 | 36 | 8.80E-11 | 2.25E-03 |
| Issac_4_1_NAQ444_J100_190802_FIXED_rere.mzML | 46930 | HCD | | 917.841 | 0 | -1.13 | 3 | K | NPDK+226.078NIVFSPLSISAALALVSLGAK | G | sp\|Q03734\|SPA3M_MOUSE | 91 | 40 | 2.78E-11 | 7.17E-04 |
| Issac_4_1_NAQ444_J100_190802_FIXED_rere.mzML | 47796 | HCD | | 879.827 | 0 | -0.28 | 3 | N | PDK+226.078NIVFSPLSISAALALVSLGAK | G | sp\|Q03734\|SPA3M_MOUSE | 134 | 78 | 4.23E-13 | 1.09E-05 |
| Issac_4_1_NAQ444_J100_190802_FIXED_rere.mzML | 20556 | HCD | | 682.349 | 0 | -0.89 | 2 | K | LK+226.078DSFLADTK | H | sp\|Q06770\|CBG_MOUSE | 68 | 53 | 1.20E-10 | 2.84E-03 |
| Issac_4_1_NAQ444_J100_190802_FIXED_rere.mzML | 10016 | HCD | | 671.350 | 0 | -1.00 | 2 | Q | K+226.078SGLQEIQGR | N | sp\|Q3KNY0\|IGFN1_MOUSE | 101 | 86 | 3.84E-11 | 9.06E-04 |
| Issac_4_1_NAQ444_J100_190802_FIXED_rere.mzML | 16450 | HCD | | 664.866 | 0 | 0.55 | 2 | R | TK+226.078TEGLVQVK | L | sp\|Q61129\|CFAI_MOUSE | 84 | 84 | 7.13E-13 | 1.68E-05 |
| Issac_4_1_NAQ444_J100_190802_FIXED_rere.mzML | 23501 | HCD | | 850.748 | 0 | 0.50 | 3 | K | K+226.078LISVDTEQSNFYLQNGPDR | I | sp\|Q61147\|CERU_MOUSE | 88 | 43 | 1.33E-11 | 3.34E-04 |
| Issac_4_1_NAQ444_J100_190802_FIXED_rere.mzML | 28301 | HCD | | 833.419 | 0 | 0.81 | 3 | R | DTANLFPHK+226.078SLTLLM(ox)NPDTK | G | sp\|Q61147\|CERU_MOUSE | 189 | 181 | 6.36E-22 | 1.60E-14 |
| Issac_4_1_NAQ444_J100_190802_FIXED_rere.mzML | 39527 | HCD | | 661.037 | 0 | -0.09 | 3 | K | TIDK+226.078PAWLGFLGPVIK | A | sp\|Q61147\|CERU_MOUSE | 88 | 78 | 2.13E-14 | 5.25E-07 |
| Issac_4_1_NAQ444_J100_190802_FIXED_rere.mzML | 39567 | HCD | | 991.553 | 1 | -1.20 | 2 | K | TIDK+226.078PAWLGFLGPVIK | A | sp\|Q61147\|CERU_MOUSE | 79 | 55 | 5.50E-12 | 1.36E-04 |
| Issac_4_1_NAQ444_J100_190802_FIXED_rere.mzML | 33769 | HCD | | 953.181 | 0 | 0.26 | 3 | A | VDLPGQQPVSEQAQQK+226.078LPLPALFK | L | sp\|Q61247\|A2AP_MOUSE | 168 | 133 | 1.51E-18 | 3.89E-11 |
| Issac_4_1_NAQ444_J100_190802_FIXED_rere.mzML | 23471 | HCD | | 678.998 | 0 | 1.08 | 3 | R | IIGGSM(ox)DAK+226.078GSFPWQAK | M | sp\|Q61646\|HPT_MOUSE | 109 | 90 | 8.51E-15 | 2.11E-07 |
| Issac_4_1_NAQ444_J100_190802_FIXED_rere.mzML | 32216 | HCD | | 834.118 | 0 | 0.95 | 3 | K | DITPTLTLYVGK+226.078NQLVEIEK | V | sp\|Q61646\|HPT_MOUSE | 113 | 73 | 6.43E-15 | 1.62E-07 |
| Issac_4_1_NAQ444_J100_190802_FIXED_rere.mzML | 8406 | HCD | | 595.296 | 0 | 1.85 | 2 | K | VNCK+226.078VTSR | F | sp\|Q61702\|ITIH1_MOUSE | 86 | 86 | 5.04E-12 | 1.16E-04 |
| Issac_4_1_NAQ444_J100_190802_FIXED_rere.mzML | 19756 | HCD | | 653.011 | 0 | 0.75 | 3 | R | NDLVSATK+226.078TQIADAKR | Y | sp\|Q61703\|ITIH2_MOUSE | 121 | 90 | 2.59E-14 | 6.39E-07 |
| Issac_4_1_NAQ444_J100_190802_FIXED_rere.mzML | 14911 | HCD | | 692.885 | 0 | 0.00 | 2 | K | AVSQGK+226.078TAGLVK | A | sp\|Q61704\|ITIH3_MOUSE | 141 | 141 | 5.25E-15 | 1.26E-07 |
| Issac_4_1_NAQ444_J100_190802_FIXED_rere.mzML | 19501 | HCD | | 501.540 | 0 | 0.55 | 4 | N | K+226.078LTEVPALVHKDTVVK | S | sp\|Q61838\|PZP_MOUSE | 146 | 113 | 7.40E-15 | 1.82E-07 |
| Issac_4_1_NAQ444_J100_190802_FIXED_rere.mzML | 20372 | HCD | | 888.955 | 0 | -0.82 | 2 | R | DLSSSDLSTASK+226.078IVK | W | sp\|Q61838\|PZP_MOUSE | 218 | 199 | 1.95E-17 | 4.78E-10 |
| Issac_4_1_NAQ444_J100_190802_FIXED_rere.mzML | 20386 | HCD | | 592.974 | 0 | 1.03 | 3 | R | DLSSSDLSTASK+226.078IVK | W | sp\|Q61838\|PZP_MOUSE | 154 | 136 | 5.97E-16 | 1.46E-08 |
| Issac_4_1_NAQ444_J100_190802_FIXED_rere.mzML | 20651 | HCD | | 888.956 | 0 | -0.62 | 2 | R | DLSSSDLSTASK+226.078IVK | W | sp\|Q61838\|PZP_MOUSE | 91 | 70 | 4.75E-13 | 1.16E-05 |
| Issac_4_1_NAQ444_J100_190802_FIXED_rere.mzML | 21746 | HCD | | 689.367 | 0 | -0.53 | 2 | K | FEVIIK+226.078M(ox)QK | T | sp\|Q61838\|PZP_MOUSE | 98 | 87 | 6.07E-11 | 1.42E-03 |
| Issac_4_1_NAQ444_J100_190802_FIXED_rere.mzML | 22026 | HCD | | 689.367 | 0 | -0.35 | 2 | K | FEVIIK+226.078M(ox)QK | T | sp\|Q61838\|PZP_MOUSE | 99 | 72 | 6.90E-11 | 1.61E-03 |
| Issac_4_1_NAQ444_J100_190802_FIXED_rere.mzML | 22408 | HCD | | 717.881 | 0 | -0.60 | 2 | K | VK+226.078ALSFYQPR | A | sp\|Q61838\|PZP_MOUSE | 131 | 127 | 1.41E-13 | 3.32E-06 |
| Issac_4_1_NAQ444_J100_190802_FIXED_rere.mzML | 22444 | HCD | | 625.685 | 0 | 0.20 | 3 | K | LTEVPALVHK+226.078DTVVK | S | sp\|Q61838\|PZP_MOUSE | 154 | 131 | 6.74E-17 | 1.65E-09 |
| Issac_4_1_NAQ444_J100_190802_FIXED_rere.mzML | 22449 | HCD | | 478.924 | 0 | 0.13 | 3 | K | VK+226.078ALSFYQPR | A | sp\|Q61838\|PZP_MOUSE | 101 | 85 | 1.16E-12 | 2.73E-05 |
| Issac_4_1_NAQ444_J100_190802_FIXED_rere.mzML | 22661 | HCD | | 717.882 | 0 | 0.34 | 2 | K | VK+226.078ALSFYQPR | A | sp\|Q61838\|PZP_MOUSE | 130 | 126 | 1.41E-13 | 3.32E-06 |
| Issac_4_1_NAQ444_J100_190802_FIXED_rere.mzML | 23360 | HCD | | 857.461 | 0 | -0.43 | 3 | K | AESPVFVQTDKPIYK+226.078PGQIVK | F | sp\|Q61838\|PZP_MOUSE | 230 | 213 | 7.16E-22 | 1.81E-14 |
| Issac_4_1_NAQ444_J100_190802_FIXED_rere.mzML | 23620 | HCD | | 857.462 | 0 | 1.49 | 3 | K | AESPVFVQTDK+226.078PIYKPGQIVK | F | sp\|Q61838\|PZP_MOUSE | 195 | 169 | 3.23E-21 | 8.19E-14 |
| Issac_4_1_NAQ444_J100_190802_FIXED_rere.mzML | 24083 | HCD | | 719.140 | 0 | 0.08 | 4 | K | AESPVFVQTDKPIYKPGQIVK+226.078FR | V | sp\|Q61838\|PZP_MOUSE | 238 | 234 | 1.67E-29 | 4.26E-22 |
| Issac_4_1_NAQ444_J100_190802_FIXED_rere.mzML | 24117 | HCD | | 958.518 | 0 | 0.64 | 3 | K | AESPVFVQTDKPIYKPGQIVK+226.078FR | V | sp\|Q61838\|PZP_MOUSE | 155 | 134 | 1.18E-22 | 3.01E-15 |
| Issac_4_1_NAQ444_J100_190802_FIXED_rere.mzML | 24265 | HCD | | 869.952 | 0 | 0.70 | 2 | K | SK+226.078AINYLISGYQR | Q | sp\|Q61838\|PZP_MOUSE | 179 | 172 | 1.42E-16 | 3.44E-09 |
| Issac_4_1_NAQ444_J100_190802_FIXED_rere.mzML | 24272 | HCD | | 580.303 | 0 | 0.63 | 3 | K | SK+226.078AINYLISGYQR | Q | sp\|Q61838\|PZP_MOUSE | 94 | 86 | 7.36E-14 | 1.78E-06 |
| Issac_4_1_NAQ444_J100_190802_FIXED_rere.mzML | 24517 | HCD | | 869.950 | 0 | -0.56 | 2 | K | SK+226.078AINYLISGYQR | Q | sp\|Q61838\|PZP_MOUSE | 170 | 166 | 1.34E-16 | 3.25E-09 |
| Issac_4_1_NAQ444_J100_190802_FIXED_rere.mzML | 24538 | HCD | | 580.303 | 0 | 0.11 | 3 | K | SK+226.078AINYLISGYQR | Q | sp\|Q61838\|PZP_MOUSE | 76 | 65 | 3.32E-13 | 8.02E-06 |
| Issac_4_1_NAQ444_J100_190802_FIXED_rere.mzML | 24779 | HCD | | 869.947 | 0 | -4.70 | 2 | K | SK+226.078AINYLISGYQR | Q | sp\|Q61838\|PZP_MOUSE | 134 | 108 | 3.10E-14 | 7.48E-07 |
| Issac_4_1_NAQ444_J100_190802_FIXED_rere.mzML | 24938 | HCD | | 689.353 | 0 | -0.71 | 3 | R | SQK+226.078EVLVTIESSGTFSK | T | sp\|Q61838\|PZP_MOUSE | 116 | 98 | 5.92E-15 | 1.47E-07 |
| Issac_4_1_NAQ444_J100_190802_FIXED_rere.mzML | 25997 | HCD | | 960.812 | 0 | -2.29 | 3 | R | LPDLPGNYVTK+226.078GSGSGCVYLQTSLK | Y | sp\|Q61838\|PZP_MOUSE | 244 | 228 | 9.92E-26 | 2.56E-18 |
| Issac_4_1_NAQ444_J100_190802_FIXED_rere.mzML | 26263 | HCD | | 960.814 | 0 | -0.06 | 3 | R | LPDLPGNYVTK+226.078GSGSGCVYLQTSLK | Y | sp\|Q61838\|PZP_MOUSE | 246 | 229 | 5.01E-27 | 1.29E-19 |
| Issac_4_1_NAQ444_J100_190802_FIXED_rere.mzML | 26571 | HCD | | 960.816 | 0 | 2.10 | 3 | R | LPDLPGNYVTK+226.078GSGSGCVYLQTSLK | Y | sp\|Q61838\|PZP_MOUSE | 194 | 159 | 3.47E-21 | 8.94E-14 |
| Issac_4_1_NAQ444_J100_190802_FIXED_rere.mzML | 26749 | HCD | | 961.146 | 1 | -2.39 | 3 | R | LPDLPGNYVTK+226.078GSGSGCVYLQTSLK | Y | sp\|Q61838\|PZP_MOUSE | 195 | 151 | 3.36E-21 | 8.65E-14 |
| Issac_4_1_NAQ444_J100_190802_FIXED_rere.mzML | 26861 | HCD | | 960.814 | 0 | 0.00 | 3 | R | LPDLPGNYVTK+226.078GSGSGCVYLQTSLK | Y | sp\|Q61838\|PZP_MOUSE | 129 | 65 | 5.73E-14 | 1.48E-06 |
| Issac_4_1_NAQ444_J100_190802_FIXED_rere.mzML | 26897 | HCD | | 813.958 | 0 | 1.12 | 4 | K | SIQIIK+226.078AESPVFVQTDKPIYKPGQIVK | F | sp\|Q61838\|PZP_MOUSE | 287 | 259 | 2.34E-26 | 6.08E-19 |
| Issac_4_1_NAQ444_J100_190802_FIXED_rere.mzML | 27101 | HCD | | 961.146 | 1 | -1.95 | 3 | R | LPDLPGNYVTK+226.078GSGSGCVYLQTSLK | Y | sp\|Q61838\|PZP_MOUSE | 148 | 121 | 3.91E-21 | 1.01E-13 |
| Issac_4_1_NAQ444_J100_190802_FIXED_rere.mzML | 29697 | HCD | | 983.515 | 0 | -1.12 | 3 | K | FTK+226.078VNTNYRPGLPFSGQVLLVDEK | G | sp\|Q61838\|PZP_MOUSE | 246 | 231 | 5.40E-27 | 1.39E-19 |
| Issac_4_1_NAQ444_J100_190802_FIXED_rere.mzML | 29712 | HCD | | 983.850 | 1 | -0.85 | 3 | K | FTK+226.078VNTNYRPGLPFSGQVLLVDEK | G | sp\|Q61838\|PZP_MOUSE | 236 | 226 | 1.41E-27 | 3.61E-20 |
| Issac_4_1_NAQ444_J100_190802_FIXED_rere.mzML | 32598 | HCD | | 1166.569 | 1 | -0.51 | 3 | K | QQNSHGGFSSTQDTVVALQALSK+226.078YGAATFTR | S | sp\|Q61838\|PZP_MOUSE | 139 | 51 | 8.10E-13 | 2.14E-05 |
| Issac_4_1_NAQ444_J100_190802_FIXED_rere.mzML | 34172 | HCD | | 568.652 | 0 | -1.18 | 3 | K | EYVLPK+226.078FEVIIK | M | sp\|Q61838\|PZP_MOUSE | 125 | 125 | 1.20E-15 | 2.89E-08 |
| Issac_4_1_NAQ444_J100_190802_FIXED_rere.mzML | 34181 | HCD | | 852.475 | 0 | -0.86 | 2 | K | EYVLPK+226.078FEVIIK | M | sp\|Q61838\|PZP_MOUSE | 161 | 140 | 2.73E-14 | 6.55E-07 |
| Issac_4_1_NAQ444_J100_190802_FIXED_rere.mzML | 34345 | HCD | | 986.516 | 0 | -0.06 | 3 | K | IKEEGTGIELTGIGSCEIANALSK+226.078LK | F | sp\|Q61838\|PZP_MOUSE | 289 | 209 | 1.22E-20 | 3.16E-13 |
| Issac_4_1_NAQ444_J100_190802_FIXED_rere.mzML | 34434 | HCD | | 852.477 | 0 | 1.65 | 2 | K | EYVLPK+226.078FEVIIK | M | sp\|Q61838\|PZP_MOUSE | 144 | 129 | 8.82E-15 | 2.12E-07 |
| Issac_4_1_NAQ444_J100_190802_FIXED_rere.mzML | 34708 | HCD | | 852.477 | 0 | 1.43 | 2 | K | EYVLPK+226.078FEVIIK | M | sp\|Q61838\|PZP_MOUSE | 115 | 85 | 4.53E-12 | 1.09E-04 |
| Issac_4_1_NAQ444_J100_190802_FIXED_rere.mzML | 38167 | HCD | | 1028.879 | 0 | 0.12 | 3 | K | YNILPVADGK+226.078APFALQVNTLPLNFDK | A | sp\|Q61838\|PZP_MOUSE | 131 | 40 | 7.25E-11 | 1.88E-03 |
| Issac_4_1_NAQ444_J100_190802_FIXED_rere.mzML | 38292 | HCD | | 1017.547 | 1 | 0.56 | 3 | K | LSPQSIYNLLPGK+226.078TVQGAFFGVPVYK | D | sp\|Q61838\|PZP_MOUSE | 179 | 133 | 1.53E-18 | 3.96E-11 |
| Issac_4_1_NAQ444_J100_190802_FIXED_rere.mzML | 38354 | HCD | | 987.889 | 0 | 0.25 | 3 | K | VK+226.078AAPLSLCALTAVDQSVLLLKPEAK | L | sp\|Q61838\|PZP_MOUSE | 270 | 260 | 6.66E-29 | 1.73E-21 |
| Issac_4_1_NAQ444_J100_190802_FIXED_rere.mzML | 38398 | HCD | | 741.169 | 0 | 0.33 | 4 | K | VK+226.078AAPLSLCALTAVDQSVLLLKPEAK | L | sp\|Q61838\|PZP_MOUSE | 202 | 128 | 4.47E-17 | 1.16E-09 |
| Issac_4_1_NAQ444_J100_190802_FIXED_rere.mzML | 38610 | HCD | | 988.222 | 1 | -1.34 | 3 | K | VK+226.078AAPLSLCALTAVDQSVLLLKPEAK | L | sp\|Q61838\|PZP_MOUSE | 142 | 97 | 2.10E-17 | 5.43E-10 |
| Issac_4_1_NAQ444_J100_190802_FIXED_rere.mzML | 38636 | HCD | | 987.889 | 0 | 0.12 | 3 | K | VK+226.078AAPLSLCALTAVDQSVLLLKPEAK | L | sp\|Q61838\|PZP_MOUSE | 140 | 87 | 9.37E-16 | 2.43E-08 |
| Issac_4_1_NAQ444_J100_190802_FIXED_rere.mzML | 39576 | HCD | | 912.167 | 0 | -0.33 | 3 | K | AAPLSLCALTAVDQSVLLLK+226.078PEAK | L | sp\|Q61838\|PZP_MOUSE | 163 | 115 | 8.72E-15 | 2.24E-07 |
| Issac_4_1_NAQ444_J100_190802_FIXED_rere.mzML | 51421 | HCD | | 960.814 | 0 | 0.51 | 3 | R | LPDLPGNYVTK+226.078GSGSGCVYLQTSLK | Y | sp\|Q61838\|PZP_MOUSE | 241 | 222 | 2.20E-25 | 5.68E-18 |
| Issac_4_1_NAQ444_J100_190802_FIXED_rere.mzML | 25518 | HCD | | 730.375 | 1 | -1.41 | 6 | R | FLPK+226.078VMLALLVSVSAEGYK+226.078K+226.078LAVWLNDMENY R | L | sp\|Q6PB70\|ANO8_MOUSE | 114 | 16 | 6.80E-11 | 1.80E-03 |
| Issac_4_1_NAQ444_J100_190802_FIXED_rere.mzML | 19843 | HCD | | 885.765 | 0 | 0.48 | 3 | R | K+226.078GGETSEM(ox)YLIQPDTSIKPYR | V | sp\|Q8K0E8\|FIBB_MOUSE | 137 | 92 | 1.27E-13 | 3.22E-06 |
| Issac_4_1_NAQ444_J100_190802_FIXED_rere.mzML | 21149 | HCD | | 810.049 | 0 | 0.38 | 3 | R | SK+226.078IQKLESDISAQM(ox)EYCR | T | sp\|Q8K0E8\|FIBB_MOUSE | 118 | 95 | 3.48E-16 | 8.68E-09 |
| Issac_4_1_NAQ444_J100_190802_FIXED_rere.mzML | 29214 | HCD | | 909.113 | 1 | -1.66 | 3 | K | KYCGLPGEYWLGNDK+226.078ISQLTR | M | sp\|Q8K0E8\|FIBB_MOUSE | 181 | 170 | 9.00E-23 | 2.28E-15 |
| Issac_4_1_NAQ444_J100_190802_FIXED_rere.mzML | 31044 | HCD | | 866.417 | 1 | 0.66 | 3 | K | YCGLPGEYWLGNDK+226.078ISQLTR | M | sp\|Q8K0E8\|FIBB_MOUSE | 116 | 103 | 5.62E-19 | 1.42E-11 |
| Issac_4_1_NAQ444_J100_190802_FIXED_rere.mzML | 17268 | HCD | | 738.897 | 0 | -0.33 | 2 | R | VSTK+226.078IQTAQFK | M | sp\|Q8K182\|CO8A_MOUSE | 118 | 114 | 6.71E-14 | 1.60E-06 |
| Issac_4_1_NAQ444_J100_190802_FIXED_rere.mzML | 23621 | HCD | | 921.817 | 0 | 0.99 | 3 | K | SHSAGVTVGVAPVK+226.078SPVSIEVTGSGSK | A | sp\|Q8K182\|CO8A_MOUSE | 228 | 204 | 9.53E-24 | 2.48E-16 |
| Issac_4_1_NAQ444_J100_190802_FIXED_rere.mzML | 10812 | HCD | | 638.803 | 0 | -0.67 | 2 | R | VGPESDK+226.078YR | L | sp\|Q8VCM7\|FIBG_MOUSE | 80 | 79 | 6.61E-13 | 1.54E-05 |
| Issac_4_1_NAQ444_J100_190802_FIXED_rere.mzML | 19715 | HCD | | 935.123 | 0 | 4.31 | 3 | K | AIQVYYNPDQPPK+226.078PGM(ox)IDSATQK | S | sp\|Q8VCM7\|FIBG_MOUSE | 115 | 53 | 2.58E-13 | 6.60E-06 |
| Issac_4_1_NAQ444_J100_190802_FIXED_rere.mzML | 20806 | HCD | | 770.391 | 0 | -0.24 | 3 | R | YLQEIYNSNNQK+226.078ITNLK | Q | sp\|Q8VCM7\|FIBG_MOUSE | 118 | 108 | 3.19E-18 | 7.91E-11 |
| Issac_4_1_NAQ444_J100_190802_FIXED_rere.mzML | 25099 | HCD | | 1096.556 | 1 | 0.13 | 3 | K | ELIK+226.078AIQVYYNPDQPPKPGM(ox)IDSATQK | S | sp\|Q8VCM7\|FIBG_MOUSE | 261 | 215 | 9.89E-23 | 2.57E-15 |
| Issac_4_1_NAQ444_J100_190802_FIXED_rere.mzML | 25114 | HCD | | 1096.221 | 0 | -0.33 | 3 | K | ELIK+226.078AIQVYYNPDQPPKPGM(ox)IDSATQK | S | sp\|Q8VCM7\|FIBG_MOUSE | 243 | 179 | 4.79E-20 | 1.25E-12 |
| Issac_4_1_NAQ444_J100_190802_FIXED_rere.mzML | 25138 | HCD | | 822.418 | 0 | 0.82 | 4 | K | ELIK+226.078AIQVYYNPDQPPKPGM(ox)IDSATQK | S | sp\|Q8VCM7\|FIBG_MOUSE | 241 | 149 | 2.35E-18 | 6.11E-11 |
| Issac_4_1_NAQ444_J100_190802_FIXED_rere.mzML | 25654 | HCD | | 602.331 | 0 | -0.41 | 3 | K | GAK+226.078ESGLYFIRPLK | A | sp\|Q8VCM7\|FIBG_MOUSE | 102 | 100 | 2.39E-14 | 5.81E-07 |
| Issac_4_1_NAQ444_J100_190802_FIXED_rere.mzML | 26282 | HCD | | 583.324 | 0 | -0.10 | 3 | K | ESGLYFIRPLK+226.078AK | Q | sp\|Q8VCM7\|FIBG_MOUSE | 143 | 120 | 6.44E-15 | 1.56E-07 |
| Issac_4_1_NAQ444_J100_190802_FIXED_rere.mzML | 31272 | HCD | | 790.040 | 0 | 0.77 | 3 | K | SSTTNGFDDGIIWATWK+226.078SR | W | sp\|Q8VCM7\|FIBG_MOUSE | 121 | 76 | 5.98E-12 | 1.50E-04 |
| Issac_4_1_NAQ444_J100_190802_FIXED_rere.mzML | 31950 | HCD | | 915.483 | 0 | -0.13 | 3 | K | KM(ox)VEEIVK+226.078YEALLLTHETSIR | Y | sp\|Q8VCM7\|FIBG_MOUSE | 190 | 140 | 2.62E-17 | 6.64E-10 |
| Issac_4_1_NAQ444_J100_190802_FIXED_rere.mzML | 31954 | HCD | | 686.865 | 0 | 0.18 | 4 | K | KM(ox)VEEIVK+226.078YEALLLTHETSIR | Y | sp\|Q8VCM7\|FIBG_MOUSE | 204 | 166 | 9.97E-18 | 2.52E-10 |
| Issac_4_1_NAQ444_J100_190802_FIXED_rere.mzML | 34203 | HCD | | 872.785 | 0 | -0.42 | 3 | K | M(ox)VEEIVK+226.078YEALLLTHETSIR | Y | sp\|Q8VCM7\|FIBG_MOUSE | 259 | 259 | 1.41E-24 | 3.55E-17 |
| Issac_4_1_NAQ444_J100_190802_FIXED_rere.mzML | 34334 | HCD | | 873.119 | 1 | -0.82 | 3 | K | M(ox)VEEIVK+226.078YEALLLTHETSIR | Y | sp\|Q8VCM7\|FIBG_MOUSE | 247 | 243 | 6.61E-24 | 1.67E-16 |
| Issac_4_1_NAQ444_J100_190802_FIXED_rere.mzML | 34480 | HCD | | 872.789 | 0 | 4.83 | 3 | K | M(ox)VEEIVK+226.078YEALLLTHETSIR | Y | sp\|Q8VCM7\|FIBG_MOUSE | 226 | 218 | 3.21E-23 | 8.10E-16 |
| Issac_4_1_NAQ444_J100_190802_FIXED_rere.mzML | 34821 | HCD | | 873.120 | 1 | 0.58 | 3 | K | M(ox)VEEIVK+226.078YEALLLTHETSIR | Y | sp\|Q8VCM7\|FIBG_MOUSE | 198 | 176 | 1.99E-22 | 5.02E-15 |
| Issac_4_1_NAQ444_J100_190802_FIXED_rere.mzML | 19687 | HCD | | 581.964 | 0 | -0.84 | 3 | K | EDK+226.078VWVYPPEKK | E | sp\|Q91X72\|HEMO_MOUSE | 88 | 59 | 8.02E-12 | 1.92E-04 |
| Issac_4_1_NAQ444_J100_190802_FIXED_rere.mzML | 24171 | HCD | | 861.401 | 0 | 0.85 | 2 | R | YYCFQGNK+226.078FLR | F | sp\|Q91X72\|HEMO_MOUSE | 145 | 145 | 1.26E-16 | 3.01E-09 |
| Issac_4_1_NAQ444_J100_190802_FIXED_rere.mzML | 24174 | HCD | | 574.603 | 0 | 0.42 | 3 | R | YYCFQGNK+226.078FLR | F | sp\|Q91X72\|HEMO_MOUSE | 128 | 128 | 1.26E-16 | 3.01E-09 |
| Issac_4_1_NAQ444_J100_190802_FIXED_rere.mzML | 24425 | HCD | | 861.400 | 0 | 0.35 | 2 | R | YYCFQGNK+226.078FLR | F | sp\|Q91X72\|HEMO_MOUSE | 151 | 151 | 1.26E-16 | 3.01E-09 |
| Issac_4_1_NAQ444_J100_190802_FIXED_rere.mzML | 24456 | HCD | | 574.937 | 1 | -0.39 | 3 | R | YYCFQGNK+226.078FLR | F | sp\|Q91X72\|HEMO_MOUSE | 84 | 81 | 2.23E-15 | 5.31E-08 |
| Issac_4_1_NAQ444_J100_190802_FIXED_rere.mzML | 24460 | HCD | | 574.603 | 0 | 0.11 | 3 | R | YYCFQGNK+226.078FLR | F | sp\|Q91X72\|HEMO_MOUSE | 71 | 70 | 6.36E-16 | 1.51E-08 |
| Issac_4_1_NAQ444_J100_190802_FIXED_rere.mzML | 27951 | HCD | | 675.859 | 0 | 1.35 | 4 | R | GPDSVFLIK+226.078EDKVWVYPPEKK | E | sp\|Q91X72\|HEMO_MOUSE | 156 | 109 | 1.55E-17 | 3.91E-10 |
| Issac_4_1_NAQ444_J100_190802_FIXED_rere.mzML | 29408 | HCD | | 865.936 | 0 | -2.68 | 2 | R | WK+226.078NPITSVDAAFR | G | sp\|Q91X72\|HEMO_MOUSE | 155 | 126 | 1.74E-15 | 4.21E-08 |
| Issac_4_1_NAQ444_J100_190802_FIXED_rere.mzML | 29426 | HCD | | 866.438 | 1 | -1.37 | 2 | R | WK+226.078NPITSVDAAFR | G | sp\|Q91X72\|HEMO_MOUSE | 130 | 130 | 6.49E-18 | 1.57E-10 |
| Issac_4_1_NAQ444_J100_190802_FIXED_rere.mzML | 32848 | HCD | | 741.854 | 0 | 0.91 | 2 | R | K+226.078WFWDFATR | T | sp\|Q91X72\|HEMO_MOUSE | 129 | 129 | 1.28E-14 | 2.99E-07 |
| Issac_4_1_NAQ444_J100_190802_FIXED_rere.mzML | 32896 | HCD | | 765.648 | 0 | 0.80 | 4 | R | WK+226.078NPITSVDAAFRGPDSVFLIKEDK | V | sp\|Q91X72\|HEMO_MOUSE | 249 | 200 | 5.83E-22 | 1.50E-14 |
| Issac_4_1_NAQ444_J100_190802_FIXED_rere.mzML | 32962 | HCD | | 765.897 | 1 | -0.54 | 4 | R | WK+226.078NPITSVDAAFRGPDSVFLIKEDK | V | sp\|Q91X72\|HEMO_MOUSE | 205 | 143 | 8.14E-17 | 2.10E-09 |
| Issac_4_1_NAQ444_J100_190802_FIXED_rere.mzML | 33100 | HCD | | 741.853 | 0 | 0.33 | 2 | R | K+226.078WFWDFATR | T | sp\|Q91X72\|HEMO_MOUSE | 127 | 127 | 1.28E-14 | 2.99E-07 |
| Issac_4_1_NAQ444_J100_190802_FIXED_rere.mzML | 33370 | HCD | | 741.853 | 0 | 0.00 | 2 | R | K+226.078WFWDFATR | T | sp\|Q91X72\|HEMO_MOUSE | 89 | 88 | 1.00E-13 | 2.34E-06 |
| Issac_4_1_NAQ444_J100_190802_FIXED_rere.mzML | 36242 | HCD | | 896.471 | 0 | -0.54 | 3 | R | WK+226.078NPITSVDAAFRGPDSVFLIK | E | sp\|Q91X72\|HEMO_MOUSE | 193 | 132 | 1.54E-17 | 3.93E-10 |
| Issac_4_1_NAQ444_J100_190802_FIXED_rere.mzML | 7107 | HCD | | 414.534 | 0 | -0.37 | 3 | K | K+226.078SCHTGLGR | S | sp\|Q921I1\|TRFE_MOUSE | 93 | 73 | 3.22E-12 | 7.52E-05 |
| Issac_4_1_NAQ444_J100_190802_FIXED_rere.mzML | 10135 | HCD | | 442.892 | 0 | 0.21 | 3 | K | VAQEHFGK+226.078GK | S | sp\|Q921I1\|TRFE_MOUSE | 107 | 103 | 2.11E-14 | 4.97E-07 |
| Issac_4_1_NAQ444_J100_190802_FIXED_rere.mzML | 12497 | HCD | | 669.315 | 0 | 0.46 | 2 | K | K+226.078TSYPDCIK | A | sp\|Q921I1\|TRFE_MOUSE | 116 | 116 | 4.41E-13 | 1.03E-05 |
| Issac_4_1_NAQ444_J100_190802_FIXED_rere.mzML | 15523 | HCD | | 568.945 | 0 | -0.32 | 3 | R | DHM(ox)K+226.078TVLPPDGPR | L | sp\|Q921I1\|TRFE_MOUSE | 98 | 47 | 1.08E-10 | 2.61E-03 |
| Issac_4_1_NAQ444_J100_190802_FIXED_rere.mzML | 15659 | HCD | | 852.914 | 0 | 0.36 | 2 | R | DHM(ox)K+226.078TVLPPDGPR | L | sp\|Q921I1\|TRFE_MOUSE | 92 | 90 | 2.79E-16 | 6.75E-09 |

| | | | | | | | | | | | | | | |
|---|---|---|---|---|---|---|---|---|---|---|---|---|---|---|
| Issac_4_1_NAQ444_J100_190802_FIXED_rere.mzML | 15778 | HCD | 568.944 | 0 | -1.39 | 3 | R | DHM(ox)K+226.078TVLPPDGPR | L | sp\|Q921I1\|TRFE_MOUSE | 106 | 80 | 8.25E-13 | 1.99E-05 |
| Issac_4_1_NAQ444_J100_190802_FIXED_rere.mzML | 16033 | HCD | 568.945 | 0 | 0.21 | 3 | R | DHM(ox)K+226.078TVLPPDGPR | L | sp\|Q921I1\|TRFE_MOUSE | 94 | 59 | 3.12E-11 | 7.53E-04 |
| Issac_4_1_NAQ444_J100_190802_FIXED_rere.mzML | 16720 | HCD | 739.654 | 0 | 0.00 | 3 | K | CAPNNK+226.078EEYNGYTGAFR | C | sp\|Q921I1\|TRFE_MOUSE | 122 | 104 | 4.32E-18 | 1.07E-10 |
| Issac_4_1_NAQ444_J100_190802_FIXED_rere.mzML | 18467 | HCD | 611.317 | 0 | -0.80 | 3 | K | K+226.078GTDFQLNQLEGKK | S | sp\|Q921I1\|TRFE_MOUSE | 134 | 119 | 5.01E-16 | 1.22E-08 |
| Issac_4_1_NAQ444_J100_190802_FIXED_rere.mzML | 19478 | HCD | 750.667 | 0 | -0.65 | 3 | K | WCAVSEHENTK+226.078CISFR | D | sp\|Q921I1\|TRFE_MOUSE | 222 | 219 | 1.48E-22 | 3.66E-15 |
| Issac_4_1_NAQ444_J100_190802_FIXED_rere.mzML | 19734 | HCD | 751.002 | 1 | -0.22 | 3 | K | WCAVSEHENTK+226.078CISFR | D | sp\|Q921I1\|TRFE_MOUSE | 176 | 117 | 1.10E-14 | 2.71E-07 |
| Issac_4_1_NAQ444_J100_190802_FIXED_rere.mzML | 19809 | HCD | 750.668 | 0 | 0.24 | 3 | K | WCAVSEHENTK+226.078CISFR | D | sp\|Q921I1\|TRFE_MOUSE | 116 | 60 | 8.00E-12 | 1.97E-04 |
| Issac_4_1_NAQ444_J100_190802_FIXED_rere.mzML | 20141 | HCD | 852.423 | 0 | -1.58 | 2 | K | K+226.078GTDFQLNQLEGK | S | sp\|Q921I1\|TRFE_MOUSE | 182 | 182 | 1.72E-17 | 4.15E-10 |
| Issac_4_1_NAQ444_J100_190802_FIXED_rere.mzML | 20191 | HCD | 568.618 | 0 | -1.40 | 3 | K | K+226.078GTDFQLNQLEGK | S | sp\|Q921I1\|TRFE_MOUSE | 86 | 59 | 6.23E-12 | 1.51E-04 |
| Issac_4_1_NAQ444_J100_190802_FIXED_rere.mzML | 20524 | HCD | 852.425 | 0 | 0.50 | 2 | K | GTDFQLNQLEGK+226.078K | S | sp\|Q921I1\|TRFE_MOUSE | 149 | 137 | 3.40E-14 | 8.21E-07 |
| Issac_4_1_NAQ444_J100_190802_FIXED_rere.mzML | 24975 | HCD | 622.326 | 0 | -0.29 | 3 | K | GK+226.078SKDFQLFSSPLGK | D | sp\|Q921I1\|TRFE_MOUSE | 134 | 131 | 4.49E-19 | 1.10E-11 |
| Issac_4_1_NAQ444_J100_190802_FIXED_rere.mzML | 25041 | HCD | 932.986 | 0 | 1.50 | 2 | K | GK+226.078SKDFQLFSSPLGK | D | sp\|Q921I1\|TRFE_MOUSE | 72 | 32 | 1.49E-11 | 3.65E-04 |
| Issac_4_1_NAQ444_J100_190802_FIXED_rere.mzML | 26511 | HCD | 1112.847 | 0 | 1.65 | 3 | R | NQQEGVCPEGSIDNSPVK+226.078WCALSHLER | T | sp\|Q921I1\|TRFE_MOUSE | 77 | 9 | 1.37E-11 | 3.55E-04 |
| Issac_4_1_NAQ444_J100_190802_FIXED_rere.mzML | 28366 | HCD | 840.427 | 0 | 0.65 | 2 | K | SK+226.078DFQLFSSPLGK | D | sp\|Q921I1\|TRFE_MOUSE | 177 | 174 | 1.17E-16 | 2.83E-09 |
| Issac_4_1_NAQ444_J100_190802_FIXED_rere.mzML | 28555 | HCD | 840.926 | 1 | -3.01 | 2 | K | SK+226.078DFQLFSSPLGK | D | sp\|Q921I1\|TRFE_MOUSE | 178 | 172 | 1.89E-16 | 4.57E-09 |
| Issac_4_1_NAQ444_J100_190802_FIXED_rere.mzML | 31782 | HCD | 621.085 | 0 | -0.79 | 4 | K | GK+226.078SKDFQLFSSPLGKDLLFK | D | sp\|Q921I1\|TRFE_MOUSE | 133 | 99 | 1.49E-16 | 3.76E-09 |
| Issac_4_1_NAQ444_J100_190802_FIXED_rere.mzML | 34718 | HCD | 642.677 | 0 | 0.38 | 3 | R | KNNGK+226.078EDLIWEILK | V | sp\|Q921I1\|TRFE_MOUSE | 125 | 112 | 1.14E-14 | 2.79E-07 |
| Issac_4_1_NAQ444_J100_190802_FIXED_rere.mzML | 34928 | HCD | 766.409 | 1 | 2.02 | 3 | K | SK+226.078DFQLFSSPLGKDLLFK | D | sp\|Q921I1\|TRFE_MOUSE | 133 | 75 | 3.05E-13 | 7.61E-06 |
| Issac_4_1_NAQ444_J100_190802_FIXED_rere.mzML | 35414 | HCD | 766.408 | 1 | 1.14 | 3 | K | SKDFQLFSSPLGK+226.078DLLFK | D | sp\|Q921I1\|TRFE_MOUSE | 165 | 152 | 9.47E-17 | 2.36E-09 |
| Issac_4_1_NAQ444_J100_190802_FIXED_rere.mzML | 35431 | HCD | 766.073 | 0 | 0.48 | 3 | K | SKDFQLFSSPLGK+226.078DLLFK | D | sp\|Q921I1\|TRFE_MOUSE | 162 | 150 | 1.10E-17 | 2.75E-10 |
| Issac_4_1_NAQ444_J100_190802_FIXED_rere.mzML | 35874 | HCD | 1004.478 | 1 | 0.20 | 3 | R | SPLEK+226.078AVSSFFSGSCVPCADPVAFPK | L | sp\|Q921I1\|TRFE_MOUSE | 193 | 141 | 1.32E-21 | 3.41E-14 |
| Issac_4_1_NAQ444_J100_190802_FIXED_rere.mzML | 35884 | HCD | 1004.143 | 0 | -0.18 | 3 | R | SPLEK+226.078AVSSFFSGSCVPCADPVAFPK | L | sp\|Q921I1\|TRFE_MOUSE | 183 | 158 | 7.39E-27 | 1.91E-19 |
| Issac_4_1_NAQ444_J100_190802_FIXED_rere.mzML | 35921 | HCD | 753.362 | 0 | 3.81 | 4 | R | SPLEK+226.078AVSSFFSGSCVPCADPVAFPK | L | sp\|Q921I1\|TRFE_MOUSE | 133 | 89 | 2.24E-17 | 5.81E-10 |
| Issac_4_1_NAQ444_J100_190802_FIXED_rere.mzML | 36149 | HCD | 1004.143 | 0 | 0.00 | 3 | R | SPLEK+226.078AVSSFFSGSCVPCADPVAFPK | L | sp\|Q921I1\|TRFE_MOUSE | 178 | 150 | 5.42E-25 | 1.40E-17 |
| Issac_4_1_NAQ444_J100_190802_FIXED_rere.mzML | 36174 | HCD | 753.359 | 0 | 0.41 | 4 | R | SPLEK+226.078AVSSFFSGSCVPCADPVAFPK | L | sp\|Q921I1\|TRFE_MOUSE | 141 | 68 | 5.23E-14 | 1.35E-06 |
| Issac_4_1_NAQ444_J100_190802_FIXED_rere.mzML | 36181 | HCD | 724.066 | 0 | 0.25 | 3 | K | DLLFK+226.078DSAFGLLRVPPR | M | sp\|Q921I1\|TRFE_MOUSE | 150 | 120 | 2.81E-14 | 6.97E-07 |
| Issac_4_1_NAQ444_J100_190802_FIXED_rere.mzML | 36426 | HCD | 860.958 | 0 | 0.14 | 2 | K | DLLFK+226.078DSAFGLLR | V | sp\|Q921I1\|TRFE_MOUSE | 167 | 166 | 6.93E-16 | 1.68E-08 |
| Issac_4_1_NAQ444_J100_190802_FIXED_rere.mzML | 36432 | HCD | 574.308 | 0 | -0.32 | 3 | K | DLLFK+226.078DSAFGLLR | V | sp\|Q921I1\|TRFE_MOUSE | 133 | 126 | 1.64E-15 | 3.96E-08 |
| Issac_4_1_NAQ444_J100_190802_FIXED_rere.mzML | 36448 | HCD | 724.066 | 0 | -0.08 | 3 | K | DLLFK+226.078DSAFGLLRVPPR | M | sp\|Q921I1\|TRFE_MOUSE | 156 | 127 | 1.59E-15 | 3.93E-08 |
| Issac_4_1_NAQ444_J100_190802_FIXED_rere.mzML | 36679 | HCD | 860.958 | 0 | -0.07 | 2 | K | DLLFK+226.078DSAFGLLR | V | sp\|Q921I1\|TRFE_MOUSE | 173 | 171 | 6.93E-16 | 1.68E-08 |
| Issac_4_1_NAQ444_J100_190802_FIXED_rere.mzML | 36702 | HCD | 724.067 | 0 | 0.51 | 3 | K | DLLFK+226.078DSAFGLLRVPPR | M | sp\|Q921I1\|TRFE_MOUSE | 146 | 88 | 3.06E-11 | 7.58E-04 |
| Issac_4_1_NAQ444_J100_190802_FIXED_rere.mzML | 36704 | HCD | 574.308 | 0 | 0.85 | 3 | K | DLLFK+226.078DSAFGLLR | V | sp\|Q921I1\|TRFE_MOUSE | 96 | 94 | 2.39E-15 | 5.78E-08 |
| Issac_4_1_NAQ444_J100_190802_FIXED_rere.mzML | 36938 | HCD | 860.958 | 0 | -0.71 | 2 | K | DLLFK+226.078DSAFGLLR | V | sp\|Q921I1\|TRFE_MOUSE | 133 | 122 | 9.06E-15 | 2.19E-07 |
| Issac_4_1_NAQ444_J100_190802_FIXED_rere.mzML | 37505 | HCD | 860.957 | 0 | -1.06 | 2 | K | DLLFK+226.078DSAFGLLR | V | sp\|Q921I1\|TRFE_MOUSE | 100 | 80 | 5.32E-13 | 1.29E-05 |
| Issac_4_1_NAQ444_J100_190802_FIXED_rere.mzML | 38013 | HCD | 860.959 | 0 | 0.71 | 2 | K | DLLFK+226.078DSAFGLLR | V | sp\|Q921I1\|TRFE_MOUSE | 80 | 72 | 3.43E-12 | 8.30E-05 |
| Issac_4_1_NAQ444_J100_190802_FIXED_rere.mzML | 38914 | HCD | 694.698 | 1 | -0.15 | 3 | K | DFQLFSSPLGK+226.078DLLFK | D | sp\|Q921I1\|TRFE_MOUSE | 60 | 37 | 3.15E-11 | 7.77E-04 |
| Issac_4_1_NAQ444_J100_190802_FIXED_rere.mzML | 47567 | HCD | 790.421 | 0 | -1.16 | 3 | R | SAGWVIPIGLLFCK+226.078LSEPR | S | sp\|Q921I1\|TRFE_MOUSE | 157 | 114 | 8.28E-15 | 2.08E-07 |
| Issac_4_1_NAQ444_J100_190802_FIXED_rere.mzML | 38761 | HCD | 885.445 | 1 | -1.56 | 3 | A | LSPTNCDASEPLAEK+226.078VLDLINK | G | sp\|Q9ESB3\|HRG_MOUSE | 171 | 120 | 5.44E-15 | 1.38E-07 |
| Issac_4_1_NAQ444_J100_190802_FIXED_rere.mzML | 38789 | HCD | 885.113 | 0 | 1.03 | 3 | A | LSPTNCDASEPLAEK+226.078VLDLINK | G | sp\|Q9ESB3\|HRG_MOUSE | 182 | 134 | 9.00E-16 | 2.29E-08 |
| Issac_4_1_NAQ444_J100_190802_FIXED_rere.mzML | 14591 | HCD | 664.355 | 0 | -0.73 | 2 | K | GTQGK+226.078IAEAVK | K | tr\|A0A0A0MQA3\|A0A0A0MQA3_MOUSE | 98 | 89 | 9.76E-13 | 2.32E-05 |
| Issac_4_1_NAQ444_J100_190802_FIXED_rere.mzML | 14612 | HCD | 664.858 | 1 | 0.78 | 2 | K | GTQGK+226.078IAEAVK | K | tr\|A0A0A0MQA3\|A0A0A0MQA3_MOUSE | 96 | 85 | 1.07E-12 | 2.55E-05 |
| Issac_4_1_NAQ444_J100_190802_FIXED_rere.mzML | 16999 | HCD | 596.982 | 0 | -0.41 | 3 | K | KVINDFVEK+226.078GTQGK | I | tr\|A0A0A0MQA3\|A0A0A0MQA3_MOUSE | 102 | 97 | 1.62E-14 | 3.95E-07 |
| Issac_4_1_NAQ444_J100_190802_FIXED_rere.mzML | 18337 | HCD | 830.923 | 0 | 1.40 | 2 | K | VINDFVEK+226.078GTQGK | I | tr\|A0A0A0MQA3\|A0A0A0MQA3_MOUSE | 159 | 156 | 2.47E-17 | 5.96E-10 |
| Issac_4_1_NAQ444_J100_190802_FIXED_rere.mzML | 20516 | HCD | 676.345 | 0 | -4.33 | 3 | K | M(ox)QHLEQTLSK+226.078ELISK | F | tr\|A0A0A0MQA3\|A0A0A0MQA3_MOUSE | 110 | 71 | 9.48E-12 | 2.32E-04 |
| Issac_4_1_NAQ444_J100_190802_FIXED_rere.mzML | 23101 | HCD | 716.381 | 0 | 0.17 | 2 | K | LVEK+226.078FLEEAK | N | tr\|A0A0A0MQA3\|A0A0A0MQA3_MOUSE | 115 | 114 | 1.07E-12 | 2.53E-05 |
| Issac_4_1_NAQ444_J100_190802_FIXED_rere.mzML | 30618 | HCD | 730.410 | 1 | -3.55 | 2 | K | ELISK+226.078FLLNR | R | tr\|A0A0A0MQA3\|A0A0A0MQA3_MOUSE | 126 | 124 | 3.37E-12 | 7.95E-05 |
| Issac_4_1_NAQ444_J100_190802_FIXED_rere.mzML | 30642 | HCD | 729.910 | 0 | -0.33 | 2 | K | ELISK+226.078FLLNR | R | tr\|A0A0A0MQA3\|A0A0A0MQA3_MOUSE | 110 | 107 | 3.37E-12 | 7.95E-05 |
| Issac_4_1_NAQ444_J100_190802_FIXED_rere.mzML | 42161 | HCD | 804.099 | 0 | -0.61 | 3 | K | KLDQDTVFALANYILFK+226.078GK | W | tr\|A0A0A0MQA3\|A0A0A0MQA3_MOUSE | 214 | 130 | 2.65E-14 | 6.64E-07 |
| Issac_4_1_NAQ444_J100_190802_FIXED_rere.mzML | 13834 | HCD | 578.314 | 0 | 1.06 | 2 | K | LALK+226.078NQDK | N | tr\|A0A0R4J0I1\|A0A0R4J0I1_MOUSE | 105 | 105 | 4.33E-11 | 9.98E-04 |
| Issac_4_1_NAQ444_J100_190802_FIXED_rere.mzML | 27336 | HCD | 609.313 | 0 | -0.70 | 3 | K | DLQILAEFHEK+226.078TR | A | tr\|A0A0R4J0I1\|A0A0R4J0I1_MOUSE | 164 | 164 | 1.67E-16 | 4.03E-09 |
| Issac_4_1_NAQ444_J100_190802_FIXED_rere.mzML | 28718 | HCD | 658.351 | 0 | -1.48 | 3 | R | K+226.078TLFSSQIEELNLPK | F | tr\|A0A0R4J0I1\|A0A0R4J0I1_MOUSE | 107 | 84 | 7.71E-14 | 1.89E-06 |
| Issac_4_1_NAQ444_J100_190802_FIXED_rere.mzML | 28735 | HCD | 658.686 | 1 | -0.34 | 3 | R | K+226.078TLFSSQIEELNLPK | F | tr\|A0A0R4J0I1\|A0A0R4J0I1_MOUSE | 116 | 98 | 3.55E-14 | 8.71E-07 |
| Issac_4_1_NAQ444_J100_190802_FIXED_rere.mzML | 38136 | HCD | 901.250 | 0 | 0.88 | 4 | K | KLSVSQVVHK+226.078AVLDVAETGTEAAAATGVIGGIRK | A | tr\|A0A0R4J0I1\|A0A0R4J0I1_MOUSE | 341 | 228 | 1.38E-23 | 3.69E-16 |
| Issac_4_1_NAQ444_J100_190802_FIXED_rere.mzML | 38140 | HCD | 721.201 | 0 | 0.76 | 5 | K | KLSVSQVVHK+226.078AVLDVAETGTEAAAATGVIGGIRK | A | tr\|A0A0R4J0I1\|A0A0R4J0I1_MOUSE | 232 | 135 | 1.22E-18 | 3.26E-11 |
| Issac_4_1_NAQ444_J100_190802_FIXED_rere.mzML | 39411 | HCD | 869.224 | 0 | -0.70 | 4 | K | KLSVSQVVHK+226.078AVLDVAETGTEAAAATGVIGGIR | K | tr\|A0A0R4J0I1\|A0A0R4J0I1_MOUSE | 354 | 114 | 2.07E-13 | 5.51E-06 |
| Issac_4_1_NAQ444_J100_190802_FIXED_rere.mzML | 39451 | HCD | 1158.966 | 1 | 0.86 | 3 | K | KLSVSQVVHK+226.078AVLDVAETGTEAAAATGVIGGIR | K | tr\|A0A0R4J0I1\|A0A0R4J0I1_MOUSE | 317 | 138 | 5.87E-16 | 1.56E-08 |
| Issac_4_1_NAQ444_J100_190802_FIXED_rere.mzML | 39457 | HCD | 1158.630 | 0 | -0.84 | 3 | K | KLSVSQVVHK+226.078AVLDVAETGTEAAAATGVIGGIR | K | tr\|A0A0R4J0I1\|A0A0R4J0I1_MOUSE | 284 | 144 | 5.92E-17 | 1.58E-09 |
| Issac_4_1_NAQ444_J100_190802_FIXED_rere.mzML | 39641 | HCD | 869.475 | 1 | -0.89 | 4 | K | KLSVSQVVHK+226.078AVLDVAETGTEAAAATGVIGGIR | K | tr\|A0A0R4J0I1\|A0A0R4J0I1_MOUSE | 156 | 72 | 3.97E-14 | 1.06E-06 |
| Issac_4_1_NAQ444_J100_190802_FIXED_rere.mzML | 43015 | HCD | 869.225 | 0 | 0.00 | 4 | K | LSVSQVVHK+226.078AVLDVAETGTEAAAATGVIGGIRK | K | tr\|A0A0R4J0I1\|A0A0R4J0I1_MOUSE | 108 | 29 | 1.22E-10 | 3.24E-03 |
| Issac_4_1_NAQ444_J100_190802_FIXED_rere.mzML | 44501 | HCD | 1116.268 | 1 | 1.00 | 3 | K | LSVSQVVHK+226.078AVLDVAETGTEAAAATGVIGGIR | K | tr\|A0A0R4J0I1\|A0A0R4J0I1_MOUSE | 185 | 86 | 3.62E-13 | 9.60E-06 |
| Issac_4_1_NAQ444_J100_190802_FIXED_rere.mzML | 44524 | HCD | 837.202 | 0 | 0.51 | 4 | K | LSVSQVVHK+226.078AVLDVAETGTEAAAATGVIGGIR | K | tr\|A0A0R4J0I1\|A0A0R4J0I1_MOUSE | 156 | 81 | 3.56E-15 | 9.45E-08 |
| Issac_4_1_NAQ444_J100_190802_FIXED_rere.mzML | 45116 | HCD | 989.883 | 0 | -0.06 | 3 | K | NQDKNIVFSPLSISAALALVSLGAK+226.078GK | T | tr\|A0A0R4J0I1\|A0A0R4J0I1_MOUSE | 220 | 131 | 1.69E-14 | 4.39E-07 |
| Issac_4_1_NAQ444_J100_190802_FIXED_rere.mzML | 45184 | HCD | 742.663 | 0 | -0.49 | 4 | K | NQDKNIVFSPLSISAALALVSLGAK+226.078GK | T | tr\|A0A0R4J0I1\|A0A0R4J0I1_MOUSE | 142 | 104 | 7.83E-17 | 2.04E-09 |
| Issac_4_1_NAQ444_J100_190802_FIXED_rere.mzML | 45202 | HCD | 742.914 | 1 | -1.05 | 4 | K | NQDKNIVFSPLSISAALALVSLGAK+226.078GK | T | tr\|A0A0R4J0I1\|A0A0R4J0I1_MOUSE | 139 | 79 | 8.97E-14 | 2.33E-06 |
| Issac_4_1_NAQ444_J100_190802_FIXED_rere.mzML | 45385 | HCD | 989.884 | 0 | 0.99 | 3 | K | NQDKNIVFSPLSISAALALVSLGAK+226.078GK | T | tr\|A0A0R4J0I1\|A0A0R4J0I1_MOUSE | 192 | 137 | 9.99E-16 | 2.60E-08 |
| Issac_4_1_NAQ444_J100_190802_FIXED_rere.mzML | 45459 | HCD | 990.216 | 1 | -1.58 | 3 | K | NQDKNIVFSPLSISAALALVSLGAK+226.078GK | T | tr\|A0A0R4J0I1\|A0A0R4J0I1_MOUSE | 172 | 132 | 3.68E-17 | 9.56E-10 |
| Issac_4_1_NAQ444_J100_190802_FIXED_rere.mzML | 45527 | HCD | 847.477 | 0 | -0.22 | 3 | Q | DK+226.078NIVFSPLSISAALALVSLGAK | G | tr\|A0A0R4J0I1\|A0A0R4J0I1_MOUSE | 160 | 104 | 1.92E-13 | 4.90E-06 |
| Issac_4_1_NAQ444_J100_190802_FIXED_rere.mzML | 45552 | HCD | 847.811 | 1 | -0.62 | 3 | Q | DK+226.078NIVFSPLSISAALALVSLGAK | G | tr\|A0A0R4J0I1\|A0A0R4J0I1_MOUSE | 131 | 43 | 8.40E-11 | 2.15E-03 |
| Issac_4_1_NAQ444_J100_190802_FIXED_rere.mzML | 46645 | HCD | 1069.944 | 0 | 0.23 | 3 | K | LALK+226.078NQDKNIVFSPLSISAALALVSLGAK | G | tr\|A0A0R4J0I1\|A0A0R4J0I1_MOUSE | 176 | 131 | 3.20E-20 | 8.39E-13 |
| Issac_4_1_NAQ444_J100_190802_FIXED_rere.mzML | 46664 | HCD | 1070.279 | 1 | 0.48 | 3 | K | LALK+226.078NQDKNIVFSPLSISAALALVSLGAK | G | tr\|A0A0R4J0I1\|A0A0R4J0I1_MOUSE | 209 | 175 | 8.42E-24 | 2.21E-16 |
| Issac_4_1_NAQ444_J100_190802_FIXED_rere.mzML | 46688 | HCD | 802.960 | 1 | -0.51 | 4 | K | LALKNQDK+226.078NIVFSPLSISAALALVSLGAK | G | tr\|A0A0R4J0I1\|A0A0R4J0I1_MOUSE | 159 | 87 | 4.56E-14 | 1.20E-06 |
| Issac_4_1_NAQ444_J100_190802_FIXED_rere.mzML | 46737 | HCD | 802.710 | 0 | 0.61 | 4 | K | LALK+226.078NQDKNIVFSPLSISAALALVSLGAK | G | tr\|A0A0R4J0I1\|A0A0R4J0I1_MOUSE | 112 | 47 | 5.95E-11 | 1.56E-03 |
| Issac_4_1_NAQ444_J100_190802_FIXED_rere.mzML | 46857 | HCD | 1008.906 | 1 | 1.23 | 3 | A | LK+226.078NQDKNIVFSPLSISAALALVSLGAK | G | tr\|A0A0R4J0I1\|A0A0R4J0I1_MOUSE | 223 | 97 | 2.55E-13 | 6.63E-06 |
| Issac_4_1_NAQ444_J100_190802_FIXED_rere.mzML | 47095 | HCD | 928.178 | 0 | 0.72 | 3 | K | NQDK+226.078NIVFSPLSISAALALVSLGAK | G | tr\|A0A0R4J0I1\|A0A0R4J0I1_MOUSE | 146 | 87 | 6.56E-14 | 1.69E-06 |
| Issac_4_1_NAQ444_J100_190802_FIXED_rere.mzML | 47377 | HCD | 1069.944 | 0 | 0.00 | 3 | K | LALKNQDK+226.078NIVFSPLSISAALALVSLGAK | G | tr\|A0A0R4J0I1\|A0A0R4J0I1_MOUSE | 201 | 153 | 2.92E-21 | 7.66E-14 |
| Issac_4_1_NAQ444_J100_190802_FIXED_rere.mzML | 47440 | HCD | 802.961 | 1 | 0.10 | 4 | K | LALK+226.078NQDKNIVFSPLSISAALALVSLGAK | G | tr\|A0A0R4J0I1\|A0A0R4J0I1_MOUSE | 130 | 45 | 1.89E-11 | 4.95E-04 |
| Issac_4_1_NAQ444_J100_190802_FIXED_rere.mzML | 47560 | HCD | 928.176 | 0 | -1.05 | 3 | K | NQDK+226.078NIVFSPLSISAALALVSLGAK | G | tr\|A0A0R4J0I1\|A0A0R4J0I1_MOUSE | 221 | 198 | 5.74E-25 | 1.48E-17 |
| Issac_4_1_NAQ444_J100_190802_FIXED_rere.mzML | 47585 | HCD | 1391.762 | 0 | -0.09 | 2 | K | NQDK+226.078NIVFSPLSISAALALVSLGAK | G | tr\|A0A0R4J0I1\|A0A0R4J0I1_MOUSE | 247 | 243 | 3.34E-27 | 8.61E-20 |
| Issac_4_1_NAQ444_J100_190802_FIXED_rere.mzML | 47592 | HCD | 696.385 | 0 | -0.09 | 4 | K | NQDK+226.078NIVFSPLSISAALALVSLGAK | G | tr\|A0A0R4J0I1\|A0A0R4J0I1_MOUSE | 194 | 177 | 2.65E-25 | 6.82E-18 |
| Issac_4_1_NAQ444_J100_190802_FIXED_rere.mzML | 47785 | HCD | 890.499 | 1 | 1.81 | 3 | N | QDK+226.078NIVFSPLSISAALALVSLGAK | G | tr\|A0A0R4J0I1\|A0A0R4J0I1_MOUSE | 205 | 157 | 2.56E-18 | 6.56E-11 |
| Issac_4_1_NAQ444_J100_190802_FIXED_rere.mzML | 47813 | HCD | 928.178 | 0 | 0.39 | 3 | K | NQDK+226.078NIVFSPLSISAALALVSLGAK | G | tr\|A0A0R4J0I1\|A0A0R4J0I1_MOUSE | 245 | 226 | 5.19E-26 | 1.34E-18 |
| Issac_4_1_NAQ444_J100_190802_FIXED_rere.mzML | 47838 | HCD | 1391.762 | 0 | -0.53 | 2 | K | NQDK+226.078NIVFSPLSISAALALVSLGAK | G | tr\|A0A0R4J0I1\|A0A0R4J0I1_MOUSE | 155 | 121 | 1.16E-16 | 2.99E-09 |
| Issac_4_1_NAQ444_J100_190802_FIXED_rere.mzML | 47864 | HCD | 1392.261 | 1 | -1.82 | 2 | K | NQDK+226.078NIVFSPLSISAALALVSLGAK | G | tr\|A0A0R4J0I1\|A0A0R4J0I1_MOUSE | 151 | 115 | 2.93E-17 | 7.55E-10 |
| Issac_4_1_NAQ444_J100_190802_FIXED_rere.mzML | 47875 | HCD | 1241.706 | 0 | -1.97 | 2 | K | NIVFSPLSISAALALVSLGAK+226.078GK | T | tr\|A0A0R4J0I1\|A0A0R4J0I1_MOUSE | 143 | 104 | 1.00E-16 | 2.56E-09 |
| Issac_4_1_NAQ444_J100_190802_FIXED_rere.mzML | 47927 | HCD | 809.135 | 0 | 0.98 | 3 | D | K+226.078NIVFSPLSISAALALVSLGAK | G | tr\|A0A0R4J0I1\|A0A0R4J0I1_MOUSE | 133 | 94 | 1.32E-15 | 3.35E-08 |
| Issac_4_1_NAQ444_J100_190802_FIXED_rere.mzML | 48065 | HCD | 928.177 | 0 | -0.07 | 3 | K | NQDK+226.078NIVFSPLSISAALALVSLGAK | G | tr\|A0A0R4J0I1\|A0A0R4J0I1_MOUSE | 222 | 199 | 3.12E-25 | 8.04E-18 |
| Issac_4_1_NAQ444_J100_190802_FIXED_rere.mzML | 48077 | HCD | 828.141 | 0 | -0.74 | 3 | K | NIVFSPLSISAALALVSLGAK+226.078GK | T | tr\|A0A0R4J0I1\|A0A0R4J0I1_MOUSE | 130 | 71 | 1.48E-12 | 3.79E-05 |
| Issac_4_1_NAQ444_J100_190802_FIXED_rere.mzML | 48096 | HCD | 828.477 | 1 | 0.98 | 3 | K | NIVFSPLSISAALALVSLGAK+226.078GK | T | tr\|A0A0R4J0I1\|A0A0R4J0I1_MOUSE | 130 | 69 | 6.77E-13 | 1.73E-05 |
| Issac_4_1_NAQ444_J100_190802_FIXED_rere.mzML | 48325 | HCD | 928.177 | 0 | -0.33 | 3 | K | NQDK+226.078NIVFSPLSISAALALVSLGAK | G | tr\|A0A0R4J0I1\|A0A0R4J0I1_MOUSE | 228 | 211 | 7.40E-26 | 1.91E-18 |
| Issac_4_1_NAQ444_J100_190802_FIXED_rere.mzML | 48441 | HCD | 928.512 | 1 | 0.75 | 3 | K | NQDK+226.078NIVFSPLSISAALALVSLGAK | G | tr\|A0A0R4J0I1\|A0A0R4J0I1_MOUSE | 189 | 162 | 7.64E-21 | 1.97E-13 |
| Issac_4_1_NAQ444_J100_190802_FIXED_rere.mzML | 48564 | HCD | 928.177 | 0 | 0.20 | 3 | K | NQDK+226.078NIVFSPLSISAALALVSLGAK | G | tr\|A0A0R4J0I1\|A0A0R4J0I1_MOUSE | 137 | 83 | 8.80E-15 | 2.27E-07 |
| Issac_4_1_NAQ444_J100_190802_FIXED_rere.mzML | 48678 | HCD | 928.511 | 1 | -0.77 | 3 | K | NQDK+226.078NIVFSPLSISAALALVSLGAK | G | tr\|A0A0R4J0I1\|A0A0R4J0I1_MOUSE | 110 | 52 | 6.24E-12 | 1.61E-04 |
| Issac_4_1_NAQ444_J100_190802_FIXED_rere.mzML | 51172 | HCD | 928.176 | 0 | -1.45 | 3 | K | NQDK+226.078NIVFSPLSISAALALVSLGAK | G | tr\|A0A0R4J0I1\|A0A0R4J0I1_MOUSE | 213 | 195 | 9.41E-26 | 2.43E-18 |
| Issac_4_1_NAQ444_J100_190802_FIXED_rere.mzML | 51449 | HCD | 928.176 | 0 | -1.38 | 3 | K | NQDK+226.078NIVFSPLSISAALALVSLGAK | G | tr\|A0A0R4J0I1\|A0A0R4J0I1_MOUSE | 165 | 74 | 1.26E-12 | 3.25E-05 |
| Issac_4_1_NAQ444_J100_190802_FIXED_rere.mzML | 31540 | HCD | 658.886 | 0 | -0.09 | 2 | K | ELISK+226.078FLLK | R | tr\|A0A0R4J0X5\|A0A0R4J0X5_MOUSE | 118 | 110 | 5.10E-11 | 1.19E-03 |
| Issac_4_1_NAQ444_J100_190802_FIXED_rere.mzML | 31816 | HCD | 658.886 | 0 | -0.09 | 2 | K | ELISK+226.078FLLK | R | tr\|A0A0R4J0X5\|A0A0R4J0X5_MOUSE | 104 | 96 | 6.36E-11 | 1.49E-03 |
| Issac_4_1_NAQ444_J100_190802_FIXED_rere.mzML | 45742 | HCD | 1106.023 | 0 | 2.10 | 2 | K | PYK+226.078CKECGK+226.078SFAKR | S | tr\|E9Q0Q3\|E9Q0Q3_MOUSE | 27 | 12 | 1.07E-10 | 2.62E-03 |
| Issac_4_1_NAQ444_J100_190802_FIXED_rere.mzML | 25272 | HCD | 766.051 | 0 | 0.40 | 3 | K | YGGSFLQGSEK+226.078SISLVQGGR | S | tr\|Q91X70\|Q91X70_MOUSE | 129 | 68 | 1.75E-11 | 4.41E-04 |
